# Plant Mobile Domain protein-DNA motif modules counteract Polycomb silencing to stabilize gene expression

**DOI:** 10.1101/2024.09.27.615353

**Authors:** Thierry Pélissier, Lucas Jarry, Margaux Olivier, Gabin Dajoux, Marie-Noëlle Pouch-Pélissier, Charles Courtois, Julie Descombin, Nathalie Picault, Guillaume Moissiard, Olivier Mathieu

## Abstract

In plants and animals, Polycomb group (PcG) proteins are crucial for development, regulating gene expression through H3K27me3 deposition and subsequent gene silencing. While Polycomb silencing target specification is increasingly understood, it remains unclear how certain genes with apparent silencing-attracting features escape this process. Here, we show that the plant mobile domain C (PMD-C) containing proteins MAINTENANCE OF MERISTEMS (MAIN), MAIN-LIKE 1 (MAIL1) and MAIL2 oppose Polycomb silencing at numerous actively transcribed genes in Arabidopsis. Mutations in *MAIN*, *MAIL1* or *MAIL2* result in PcG-dependent ectopic H3K27me3 deposition, often associated with transcriptional repression. We show that MAIL1, which functions in concert with MAIN, and MAIL2 target distinct gene sets and associate with chromatin at specific DNA sequence motifs. We demonstrate that the integrity of these motif sequences is essential for promoting expression and antagonizing H3K27me3 deposition. Our results unveil a novel system opposing Polycomb silencing involving PMD-C protein-DNA motif modules, expanding our understanding of eukaryotic gene regulation mechanisms.

## Main

Chromatin-based modulation of gene expression is mediated by enzymatic complexes that catalyze post-translational histone modifications and alter chromatin structure. The evolutionarily conserved Polycomb group (PcG) proteins act as a cellular memory system, mediating epigenetic gene repression. PcG proteins form multi-subunit complexes, primarily Polycomb Repressive Complex 1 (PRC1) and Polycomb Repressive Complex 2 (PRC2), which often work cooperatively to establish and maintain gene repression. PRC2 catalyzes the trimethylation of lysine 27 on histone H3 (H3K27me3), a hallmark generally associated with repressed chromatin, while PRC1 mediates ubiquitination on lysine 121 of H2A (H2AK121ub) in plants (or lysine 119 in animals). In the *Arabidopsis thaliana* genome, these two histone marks frequently colocalize, with PRC1-mediated H2AK121ub being required for the incorporation of H3K27me3 at most genes targeted by PcG-mediated silencing^1^.

The dynamic regulation of gene expression necessitates mechanisms to counteract PcG-mediated regulation. In *Arabidopsis*, several factors have been identified that oppose PcG activity. These include several histone methyltransferases of the ARABIDOPSIS TRITHORAX (ATX) and ARABIDOPSIS TRITHORAX RELATED (ATXR) protein families, which catalyze H3K4 methylation, thereby promoting active chromatin states^2–5^. Another important group of proteins counteracting PcG-mediated silencing are H3K27me3 demethylases. In Arabidopsis, these include the Jumonji C domain-containing histone demethylases EARLY FLOWERING 6 (ELF6), RELATIVE OF EARLY FLOWERING 6 (REF6), and JMJ13^6–9^. Additionally, ANTAGONIST OF LIKE HETEROCHROMATIN PROTEIN 1 (ALP1) and ALP2 have been proposed to interfere with PcG silencing at a small subset of PcG targets by associating with PRC2 and potentially inhibiting its activity^10,11^, although mutations in either *ALP* genes do not alter H3K27me3 levels.

We previously reported that two plant-specific proteins, MAINTENANCE OF MERISTEMS (MAIN) and MAIN-LIKE 1 (MAIL1), are important factors for plant development^12–14^. These proteins are characterized by a Plant Mobile Domain of type C (PMD-C) and have been implicated in the regulation of transposable elements and gene expression. However, their precise molecular functions and potential roles in epigenetic regulation remain poorly understood. Here we demonstrate that MAIN, MAIL1 and the homologous protein MAIL2 counteract PcG-mediated silencing at specific genomic loci by preventing ectopic H3K27me3 incorporation.

## Results

### Mutations in *NDX* and PRC core components alleviate *mail1*-associated phenotypes

Mutations in components of the MAIL1/MAIN/PP7L complex result in severe developmental defects and transcriptional abnormalities characterized by loss of silencing at several TEs and mis-expression of numerous protein-coding genes^12–15^. To uncover the molecular basis of these defects, we conducted a genetic screen for suppressors of the *mail1* mutant phenotypes. Screening individuals from an ethyl methanesulfonate (EMS) mutagenized population of *mail1* containing the *L5-GUS* transgene^16^, we identified a mutant line exhibiting both improved plant development and suppression of L5 silencing release (Fig. 1a,b). Compared to *mail1* plants, the suppressed plants showed faster growth and greener leaves. The traits followed a 1/3 ratio in the F2 progeny from a backcross, indicating a single-locus nuclear recessive causative mutation. Mapping-by-sequencing using an outcross F2 population identified a knock-out allele of *AT4G03090* (*NODULIN HOMEOBOX; NDX*) that we named *ndx-7*, associated with a C to T mutation leading to a premature stop codon (Supplementary Fig. 1a). Crossing *mail1* plants with an independent *ndx-4* T-DNA insertion mutant line and introduction of an *NDX* genomic copy in the *mail1 ndx-7* background confirmed that mutation of *NDX* alleviated *mail1*-associated developmental and silencing defects (Supplementary Fig. 1b-d). Beyond the *L5-GUS* transgene, RT-qPCR analysis revealed strong suppression of *mail1*-transcriptional activation of several endogenous targets in *mail1 ndx-7* plants (Fig. 1c). Consistent with MAIL1’s role in a complex with MAIN, introducing *ndx-7* in the *main* background also mitigated *main*-associated developmental defects and GUS upregulation (Supplementary Fig. 1b,d).

**Fig. 1.**
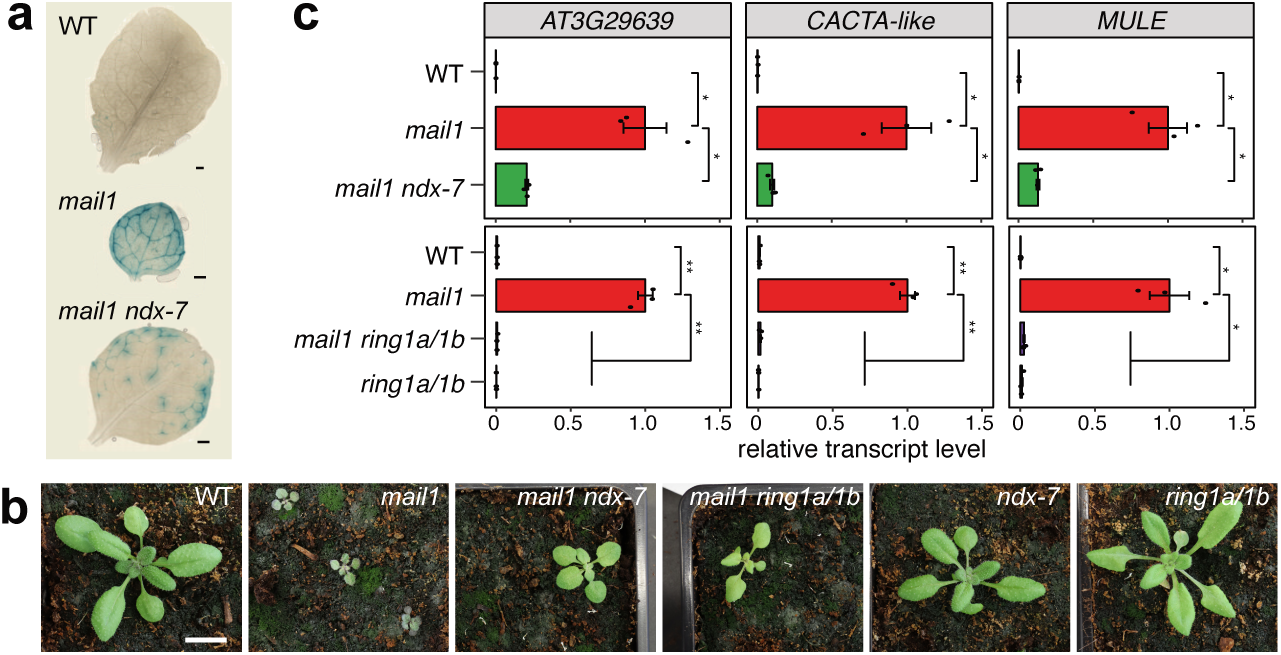
NDX and PRC1 RING1A/1B components suppress *mail1*-induced defects. (**a**) Histochemical staining for GUS activity in leaves from plants of the indicated genotypes. Scale bar is 500 µm. (**b**) Developmental phenotype of three-week-old WT, *mail1*, *mail1 ndx-7*, *mail1 ring1a/1b*, *ndx-7* and *ring1a/1b* seedlings. Scale bar is 1 cm. (**c**) Expression analysis of *mail1*-overexpressed loci by quantitative RT–PCR in the indicated genotypes. Transcript levels are normalized to *ACT2* and further normalized to *mail1*; values represent means from three biological replicates ± s.e.m. Asterisks mark statistically significant differences from the *mail1* mutant (unpaired, two-sided Student’s t-test, * *P* < 0.05; ** *P* < 0.01).

NDX is a transcription factor recently found to directly interact with PRC1 core components RING1A and RING1B, regulating a set of ABA-responsive genes^17,18^. Introgression of the *mail1* mutation into a *ring1a ring1b* (*ring1a/1b*) double mutant backgound^18^ resulted in a similar reversal of *mail1*-associated phenotypic and transcription defects (Fig. 1b,c). NDX can also indirectly interact with the transcription factor VIVIPAROUS1/ABI3-LIKE (VAL1), which is involved in PRC1 recruitment at the *FLC* gene^19–21^, through its association with RING1A and RING1B. Crossing *val1* plants, which appear indistinguishable from WT, with *mail1* or *main* mutant lines resulted in double mutants with very severe developmental phenotype characterized by the apparent absence of functional shoot apical meristem, and unable to initiate post-embryonic differentiation (Supplementary Fig. 1e). This indicates that *ndx* and *ring1a/1b* suppression of *mail1* is independent of *VAL1*.

To assess the genome-wide importance of *NDX* and *RING1A/1B* on *mail1-*associated transcriptional defects, we performed RNA-seq experiments using WT, *ndx*, *ring1a/1b*, *mail1*, *mail1 ndx* and *mail1 ring1a/1b* lines. To account for the variability we observed in *mail1* plant phenotypic severity -possibly due to subtle differences in growth conditions-we used four RNA-seq replicates to define a robust set of *mail1*-specific misregulated targets, including differentially expressed genes and a subset of upregulated TEs (Fig. 2a and Supplementary Fig. 2a). Gene ontology (GO) analysis revealed that 29% of *mail1*-upregulated genes (79 out of 273) are linked to hormone and stress pathways (Supplementary Fig. 2b), consistent with *mail1* enhancing plant sensitivity to environmental cues. The expression of most misregulated loci was unaffected in *ndx* or *ring1a/1b* mutants (Fig. 2a). Combining either *ndx* or *ring1a/1b* mutations with *mail1* resulted in an overall suppression of TE upregulation (Fig. 2a,b). Similarly, expression levels of *mail1*-upregulated genes tend to significantly return towards WT levels in *mail1 ndx* and *mail1 ring1a/1b* (Fig. 2a,b). Additionally, *mail1-*associated downregulation was suppressed for about 65% of the genes (62 to 68/95) in both mutant combinations (Fig. 2a,b and Supplementary Fig. 2c). These results indicate that *NDX* and *RING1A/1B* are required for the transcriptional misregulation induced in the absence of *MAIL1*.

**Fig. 2.**
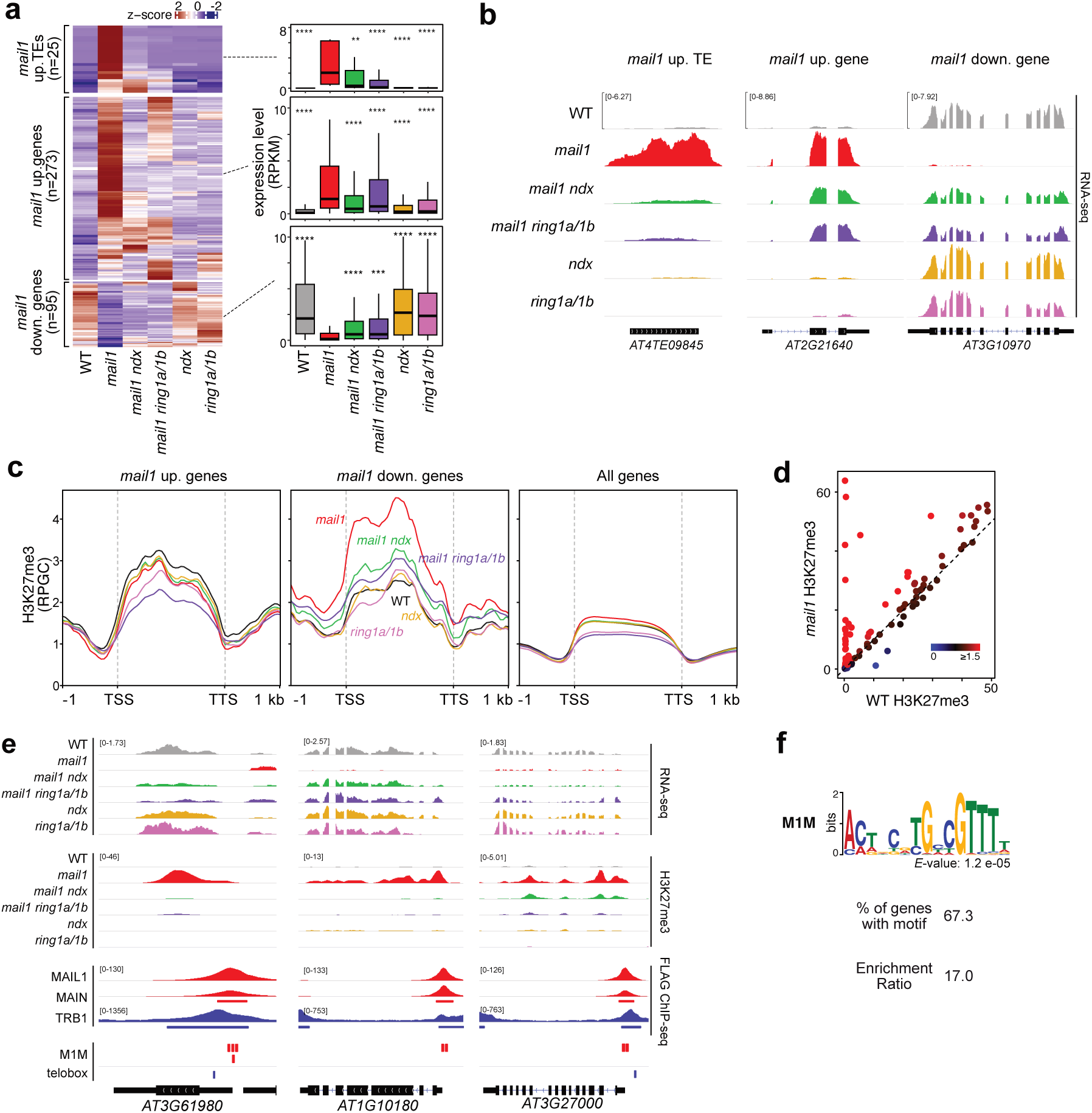
*mail1*-associated TE and gene misregulation is dependent on NDX or RING1A/1B function. (**a**) Heatmaps of RNA-seq expression z-score computed for *mail1* upregulated TEs and misregulated genes across the indicated genotypes (left). Means of two biological replicates were computed. Boxplots with the same data expressed in RPKM are shown on the right. Center lines show the medians; box limits indicate the 25th and 75th percentiles; whiskers extend 1.5 times the interquartile range from the 25th and 75th percentiles. Asterisks mark statistically significant differences from the *mail1* mutant (Wilcoxon rank sum test with continuity correction; two-sided, ** *P* < 0.01; \*\*\**P* < 0.001; \*\*\*\**P* < 0.0001). (**b**) Genome browser tracks showing mRNA profiles (CPM) in the indicated genotypes at *mail1*-misregulated loci. (**c**) Metagene plots of H3K27me3 accumulation (RPGC) in the indicated genotypes at *mail1*-upregulated, *mail1*-downregulated genes compared to all Arabidopsis genes. One of the two biological replicates is shown (**d**) H3K27me3 level variations in *mail1* compared to WT at *mail1*-downregulated genes. (**e**) Representative genome browser views of RNA-seq (CPM) and H3K27me3 ChIP-seq (RPGC) profiles in the indicated genotypes at H3K27me3-enriched genes in *mail1*. MAIL1-FLAG and MAIN-FLAG associated patterns are shown together with TRB1-FLAG data (GEO-GSM6895939 retrieved from^30^). Predicted Mail1 (M1M) and telobox motifs are indicated. (**f**) MEME prediction of a differentially enriched DNA motif within the 1kb region surrounding TSS of H3K27me3-enriched genes in *mail1*. A control promoter set was randomly selected from genes with similar H3K27me3 and expression patterns in wild type plants (ranging from 0 to 3 RPKM and 0.1 to 15 RPKM, respectively). Percentage of regions exhibiting the M1M motif and relative enrichment ratio are indicated.

To determine the impact of mutations in the PRC2 core component CLF or SWN, we generated *mail1 clf-*29 and *mail1 swn*-7 double mutants and performed RNA-seq. CLF and SWN act largely redundantly to catalyze H3K27me3 deposition in vegetative tissues, although certain loci appear to be preferentially targeted by one or other H3K27 methyltransferases^22^. We found that *mail1*-induced transcriptional misregulation at genes was significantly suppressed in *mail1 clf* but not in *mail1 swn* (Supplementary Fig. 2d), although both double mutants showed developmental defects mostly similar to those of *mail1* (Supplementary Fig. 2e). This indicates that mutation of *CLF* tends to suppress *mail1*-induced transcriptional defects, albeit to an insufficient extent to result in an improved phenotype, most likely because of the functional redundancy with SWN^22^.

Our results show that *NDX, RING1A/1B,* and to a lesser extent *CLF*, are required for the transcriptional misregulation induced in the absence of *MAIL1,* suggesting a potential link between MAIL1 and PcG-mediated silencing.

### MORC1 and MORC2 dampen TE activation in *mail1* suppressor backgrounds

MICRORCHIDIA (MORC) proteins are ATPases required for TE silencing and heterochromatin condensation^23^. We recently linked silencing defects at a subset of TEs to downregulation of MORC1 in *mail1* plants^15^. In both suppressor backgrounds, loss of TE activation correlated with increased *MORC1* transcript accumulation relative to *mail1*, albeit limited in *mail1 ndx* plants (Supplementary Fig. 3a). Interestingly, expression of the closely related *MORC2* gene, also involved in TE silencing^24^, showed a four to six fold induction in the presence of *ndx* or *ring1a/1b* mutations (Supplementary Fig. 3a). We tested whether *MORC2* could contribute to the reestablishment of TE silencing by introgressing the *morc2-1* mutation into the *mail1 ndx* background and performing RNA-seq to monitor TE expression. Of the 25 *mail1*-upregulated TEs, at least six showed dependency towards MORC2 for their repression in *mail1 ndx* plants (Supplementary Fig. 3b,c), and we previously showed that overexpressing *MORC1* in the *mail1* background also repressed some *mail1*-activated TEs^15^ (Supplementary Fig. 3d). This is consistent with MORC1 and MORC2 acting redundantly in TE silencing^24^ and suggests that upregulation of both *MORC1* and *MORC2* contributes to dampening TE activation in *mail1 ndx* and *mail1 ring1a/1b*.

### Gene silencing in *mail1* correlates with ectopic H3K27me3 incorporation

In light of the suppressor effect of *ndx* and *ring1a/1b* mutations and the known interaction of NDX and RING1A/1B in PcG-mediated silencing^17,19^, we explored a possible link between *mail1*-associated gene misregulation and the PcG-mediated silencing diagnostic mark H3K27me3. Chromatin immunoprecipitation sequencing (ChIP-seq) analyses revealed no overall change in H3K27me3 levels in *mail1* when considering all Arabidopsis genes (Fig. 2c). Likewise, the *ndx* mutation did not significantly alter H3K27me3 distribution over genes or TEs (Fig. 2c and Supplementary Fig. 4). In *mail1,* upregulated genes showed slight loss of H3K27me3. The addition of the *ndx* mutation had no additional effect, while H3K27me3 levels were further reduced at these genes in *mail1 ring1a/1b* mutants (Fig. 2c and Supplementary Fig. 5a). Despite this reduction, both *ndx* and *ring1a/1b* mutations tend to suppress *mail1*-

induced transcriptional activation of these genes (Fig. 2a), suggesting that their misregulation in *mail1* is not directly connected to H3K27me3 variation. Strikingly, gene downregulation in *mail1* correlated with prominent H3K27me3 enrichment, which was largely reversed upon additional *NDX* or *RING1A/1B* depletions (Fig. 2c and Supplementary Fig. 5a) that also tended to restore expression of these genes (Fig. 2a). Many genes acquired high levels of H3K27me3 *de novo* in *mail1* (Fig. 2d), correlating with transcriptional repression (Supplementary Fig. 5b), and this was largely reversed in the *mail1 ndx* and *mail1 ring1a/1b* suppressed lines (Fig. 2e and Supplementary Fig. 5c). One such gene was *MORC1*, gaining H3K27me3 in *mail1*, which was completely lost in *mail1 ring1a/1b* (Supplementary Fig. 5c). We also found that its neighbor gene *MORC2* was subject to PcG-mediated silencing in the WT, explaining its upregulation in the presence of *ndx* or *ring1a/1b* mutations (Supplementary Fig. 3a,5c). Interestingly, H3K27me3-enriched genes in *mail1* included *SPO11-1* (Supplementary Fig. 5c), which is mostly expressed in flowers and encodes a topoisomerase-like protein essential for meiotic recombination and fertility^25,26^. Gain of H3K27me3 was maintained in *mail1* immature flowers, correlating with about a 70% decrease in *SPO11-1* transcript accumulation (Supplementary Fig. 5d). Such level of *SPO11-1* downregulation was reported to significantly affect gamete viability^26^, and thus likely contributes to *mail1* hypofertility^27^.

RING1A and RING1B are largely required for PRC1-mediated H2A ubiquitination^18^, playing a key role in recruiting H3K27me3 at numerous loci^1^. This is likely the case at genes gaining H3K27me3 in *mail1* (Supplementary Table 1) as these showed concomitant enrichment in H2AK121ub (Supplementary Fig. 6a,b). Together, these results suggest that loss of *MAIL1* directly leads to PcG-mediated silencing at many genes.

To search for possible common features associated with the gain of H3K27me3 at genes in *mail1*, we performed a DNA motif-based sequence analysis on the 1-kb region surrounding the transcription start site (TSS) of these genes. Four conserved DNA motifs were detected in 60-70% of the genes (Supplementary Fig. 6c), but only the top-scoring motif was significantly enriched compared to the control set (Fig. 2f). We named this motif M1M (for MAIL1/MAIN Motif), which notably extends a previously reported “down” motif partially enriched among *mail1-* and *main*-downregulated genes^13^. Interestingly, the other motifs included telobox and GA-repeat elements, two known polycomb response elements (PREs) bound by transcription factors to tether PRC2 complexes to specific genes^22,28,29^. Using previously published data^30^, we found that the TELOMERE-REPEAT BINDING FACTOR (TRB)1, TRB2 and TRB3, which bind telobox elements were loaded in wild-type plants close to the TSS of a significant fraction (31/49) of *mail1* H3K27me3-enriched genes (Fig. 2e and Supplementary Fig. 6d,e). Together, these results suggest that the MAIL1/MAIN complex might promote the expression of some genes by preventing, directly or indirectly, PcG-mediated silencing. Upon MAIL1/MAIN depletion, PRC2 recruitment, through TRB proteins or other factors, requires PRC1 activity to be robustly established, potentially through a positive feedback loop^1^.

### MAIL1/MAIN complexes are specifically targeted to M1M motifs

To explore the genomic localizations of MAIL1 and MAIN *in vivo*, we performed ChIP-seq using transgenic lines expressing functional FLAG- or MYC-tagged versions of MAIL1 or MAIN under their native promoters in their respective mutant backgrounds^13^. MAIL1 and MAIN largely colocalized and were predominantly enriched at proximal promoter regions of genes (Fig. 3a-d). Notably, they bound approximately 68% of the genes that gain H3K27me3 in *mail1*, but showed no enrichment at genes with reduced H3K27me3 levels (Fig. 3e and Supplementary Fig. 7a). These H3K27me3-enriched genes showed similar transcriptional downregulation and H3K27me3 accumulation in *mail1*, *main*, and *mail1 main* mutants, consistent with MAIL1 and MAIN acting in the same regulatory complex (Fig. 3f). Importantly, motif analysis revealed that approximately 86% of MAIL1/MAIN-bound regions harbor the M1M motif (Supplementary Fig. 7b), strongly suggesting a functional role for M1M in recruiting MAIL1/MAIN to proximal promoter regions (Fig. 2e, 3e).

**Fig. 3.**
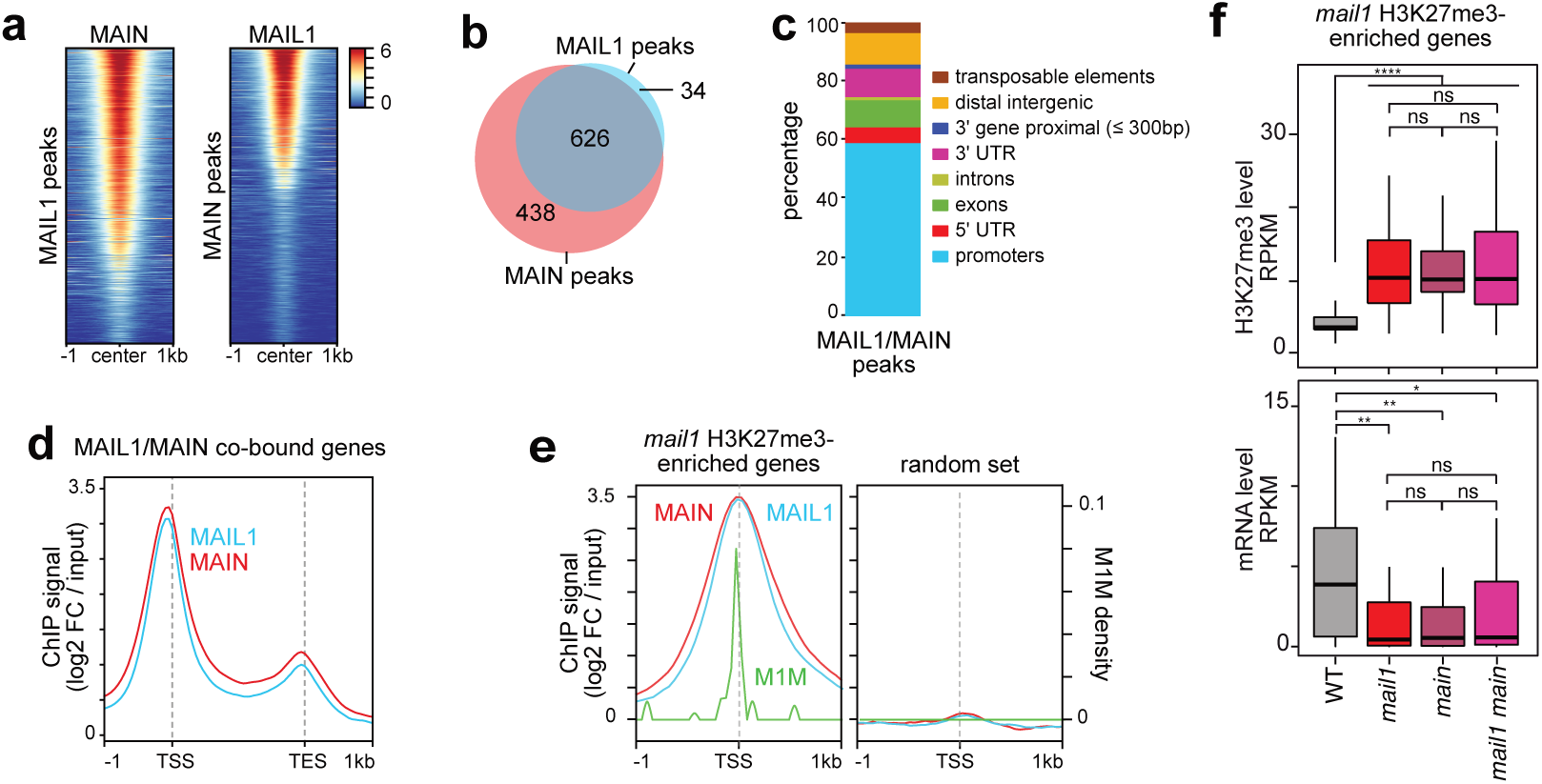
MAIL1 and MAIN are mainly co-targeted to M1M at gene proximal promoters. (**a**) Heatmaps showing MAIN and MAIL1 ChIPseq signals (Log2 FC / input) over MAIL1 (n = 660) and MAIN (n = 1064) peaks respectively. Peaks were aligned at their center position. (**b**) Overlap between MAIL1 and MAIN peaks. (**c**) Diagram illustrating the genomic distribution of MAIL1/MAIN common peaks (n = 626). (**d**) Metagene plot showing MAIL1 and MAIN enrichment at MAIL1/MAIN co-bound genes (n = 670). Co-bound genes displayed MAIL1/MAIN peak overlapping over the 200 bp region upstream TSS (proximal promoter targeted genes) or encompassed at least 50% of the peak within the TSS - TES region (body targeted genes). (**e**) Metaplots showing MAIL1, MAIN and M1M enrichment at promoter region of *mail1*-H3K27me3 enriched genes (n=49; left panel). An equivalent random set of genes is shown as control (right panel). (**f**) transcript and H3K27me3 levels, expressed in RPKM, at *mail1*-H3K27me3 enriched genes in the indicated genotypes. Center lines show the medians; box limits indicate the 25th and 75th percentiles; whiskers extend 1.5 times the interquartile range from the 25th and 75th percentiles. The effect of genotype was verified with a Wilcoxon rank sum test with continuity correction; two-sided (ns. not significant; * *P* < 0.05; ** *P* < 0.01; *** *P* < 0.001; **** *P* < 0.0001).

### NDX and RING1A are enriched at MAIL1/MAIN binding regions

To further explore how mutations of NDX may suppress *mail1*, we sought to determine the genomic localization of NDX. We generated transgenic lines expressing functional FLAG- or MYC-tagged NDX in *ndx-7* and *mail1 ndx* mutant backgrounds (Supplementary Fig. 8a,b). However, ChiP-seq experiments using anti-MYC or anti-FLAG antibodies were unsuccessful, potentially due to low NDX protein abundance. Quality control analysis indicated that recently published NDX ChIP-seq data^31^ were not suitable for further use. We therefore turned to a CUT&Tag approach, which provided good quality data only for the *ndx-7* background. Both NDX-MYC and FLAG-NDX showed similar genomic distributions, with clear enrichment at genes, particularly in promoter regions upstream of the TSSs, and depletion at TEs, especially those located in pericentromeric heterochromatin (Supplementary Fig. 8c,d). This distribution is consistent with NDX belonging to a large family of transcription factors^32,33^. Furthermore, re-analysis of published RING1A ChIP-seq data^34^ revealed substantial colocalization with NDX, consistent with their previously reported physical interaction^17,19^ (Supplementary Fig. 8e,f). Notably, both NDX and RING1A were enriched at MAIL1/MAIN binding sites (Supplementary Fig. 8e), suggesting that MAIL1/MAIN may function to prevent the recruitment of downstream PcG-silencing components at their target genes.

### MAIL2 function is essential to plant development and opposes PcG-mediated silencing at distinct loci

We previously reported that *MAIL1* and *MAIN* are Brassicaceae-specific paralogues of *MAIL2*, which is widely distributed among plant species^13^. The *MAIL2* gene appears to be under strong purifying selection^13^ suggesting that it may fulfill an important function. In line with this assumption, the *mail2-1* transfer DNA mutant displayed a very severe developmental phenotype. The shoot apical meristem (SAM) was unable to initiate a proper post-embryonic program, with plants rarely developing true leaves and typically arresting growth at the two cotyledons stage (Fig. 4a and Supplementary Fig. 9a,b). Notably, we were unable to obtain homozygous mutants in the progeny segregating from a self-fertilized heterozygous *mail2* CRISPR mutant, suggesting that *mail2-1* may not be a null mutant.

**Fig. 4.**
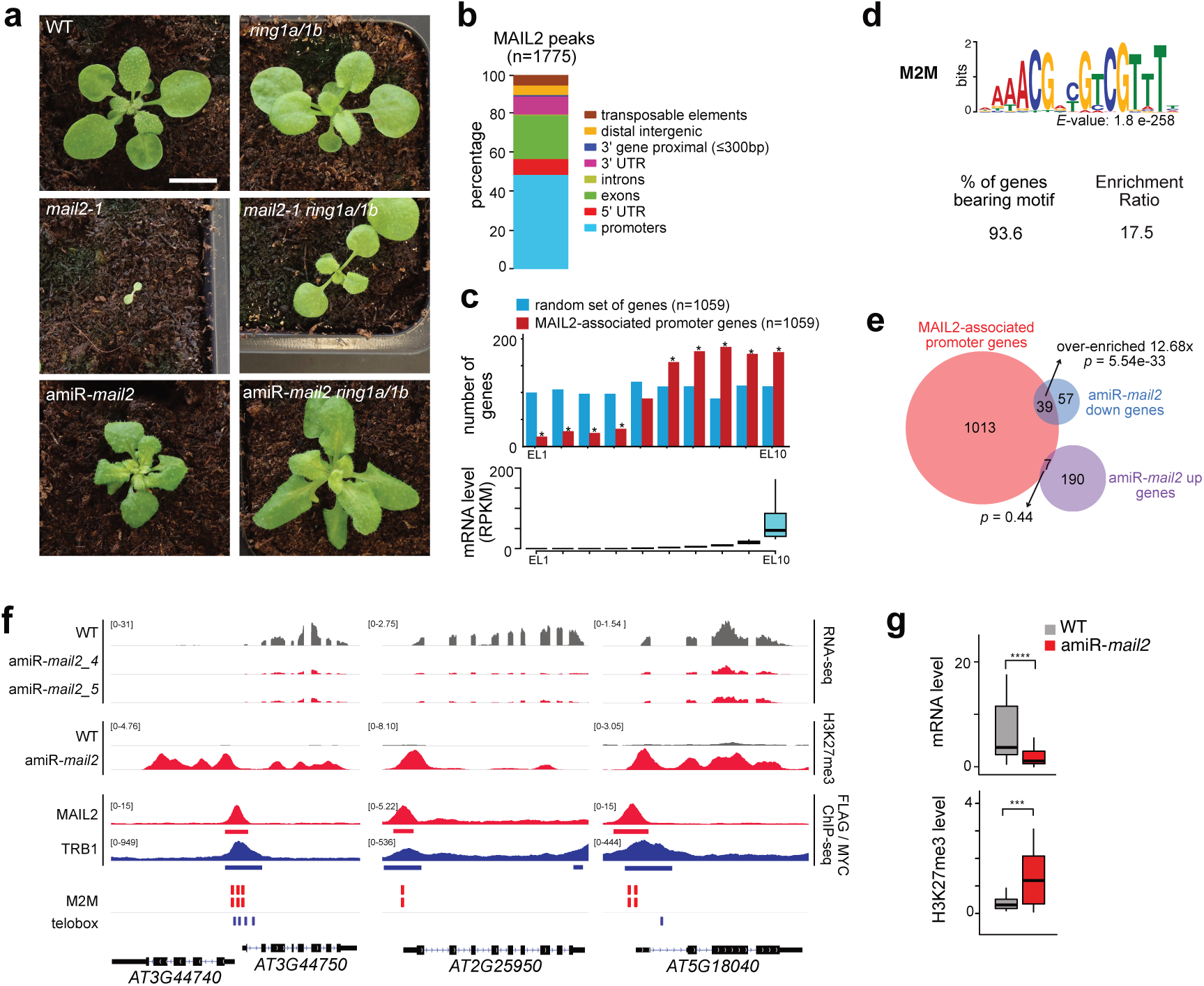
MAIL2 mainly associates to gene promoters and protects numerous actively transcribed genes from H3K27me3-mediated PcG silencing. (**a**) Photos of three-week-old seedlings of indicated genotype. Scale bar is 1 cm. (**b**) Genomic features associated with MAIL2 peaks. (**c**) Distribution of MAIL2-associated promoter genes in relation to their expression level in WT plants. A random set of the same number of genes is shown for comparison. Asterisks mark statistically significant differences from an equal representation in each decile (Fisher’s exact test, *P*-value < 1e-04). All Arabidopsis genes were split in ten deciles based on their WT expression level (RPKM; bottom diagram). Center lines show the medians; box limits indicate the 25th and 75th percentiles; whiskers extend 1.5 times the interquartile range from the 25th and 75th percentiles. (**d**) Identification of a highly enriched motif within the 1kb region surrounding TSS of genes harboring MAIL2 peak at proximal promoter (n = 1059). Analysis was carried out using MEME in differential enrichment mode and an equivalent control set randomly selected among genes ranging in the five deciles with the highest expression level in the WT. Percentage of regions exhibiting the M2M motif and relative enrichment ratio are indicated. (**e**) Overlap between MAIL2-associated promoter genes and misregulated genes in amiR-*mail2* lines. Fold change enrichment compared to random distribution and p-value from hypergeometric tests are indicated. (**f**) Representative genome browser views of RNA-seq (CPM) and H3K27me3 ChIP-seq (RPGC) profiles in the indicated genotypes at MAIL2-associated promoter genes downregulated in amiR*-mail2* background. MAIL2-MYC associated patterns are shown together with TRB1-FLAG data (GEO-GSM6895939 retrieved from^30^) and predicted MAIL2 (M2M) and telobox motif positions. (**g**) Box plots showing transcript and H3K27me3 levels (RPKM) in WT and amiR-*mail2* lines at MAIL2-associated promoter genes downregulated (n = 39) in hypomorphic amiR-*mail2* background. Center lines show the medians; box limits indicate the 25th and 75th percentiles; whiskers extend 1.5 times the interquartile range from the 25th and 75th percentiles. The effect of genotype was verified with a Wilcoxon rank sum test with continuity correction; two-sided (ns. not significant; *** *P* < 0.001; **** *P* < 0.0001).

The MAIL1, MAIN, and MAIL2 proteins share a highly homologous PMD-C domain^27^ (Supplementary Fig. 10), which is the only identified domain in these proteins. Given the similarity in structure, we hypothesized that MAIL2 might be involved in a protection mechanism against polycomb silencing similar to the one identified for MAIL1/MAIN. To test this hypothesis, we introduced *ndx*, *clf*, or *ring1a/1b* mutations into the *mail2-1* background through crossing. The resulting mutants showed varying degrees of phenotypic suppression: *mail2-1 ndx* and *mail2-1 clf* double mutants developed a first pair of leaves before arresting growth, while *mail2-1 ring1a/1b* mutants displayed more extensive development, progressing through the rosette stage and approaching the floral transition (Fig. 4a and Supplementary Fig. 9a). These observations indicate that *ndx*, *clf*, and *ring1a/1b* mutations partially suppress the *mail2-1* developmental phenotype, mirroring their effects on *mail1* and *main* mutants. This suggests a functional similarity between MAIL2 and MAIL1/MAIN in counteracting PcG-mediated silencing, implying that this anti-PcG silencing function may be an ancestral feature of the PMD-C protein family.

We determined the genomic localization of MAIL2 by ChIP-seq using a transgenic line expressing a functional, tagged version of MAIL2 (Supplementary Fig. 9c). This analysis identified 1775 MAIL2-enriched regions across the genome, largely distinct from those bound by MAIL1/MAIN (Supplementary Fig. 11a), yet similarly associated with elevated NDX and RING1A occupancy (Supplementary Fig. 8e). As in the case of MAIL1 and MAIN, these regions were predominantly associated with genes, particularly within proximal promoter regions (1059 genes) (Fig. 4b and Supplementary Fig. 11b). These genes were significantly overrepresented in the five deciles showing the highest expression level in the WT (Fig. 4c).

We performed DNA motif-based sequence analyses using MAIL2 peaks or the 1kb region surrounding TSS of MAIL2-associated gene promoters and identified a conserved sequence element, which we named MAIL2-Motif (M2M), present in nearly 95% of both the peaks and promoter regions (Fig. 4d and Supplementary Fig. 11c-e). Intriguingly, the M2M consensus resembles that of the M1M motif but in a palindromic configuration, suggesting M1M may have derived from M2M (Supplementary Fig. 11f), and mirroring the evolutionary relationship between the *MAIL1/MAIN* genes and *MAIL2*. As observed for MAIL1/MAIN-target genes, GAA-repeats and telobox elements were also present in a substantial fraction of MAIL2-associated gene promoters (Supplementary Fig. 11c).

To facilitate exploration of MAIL2 function, we generated hypomorphic *mail2* mutants using artificial microRNAs (amiRNAs) targeting *MAIL2* mRNA (amiR-*mail2*) (Fig. 4a and Supplementary Fig. 9b). While many amiR-*mail2* plants displayed severely compromised SAM development under *in vitro* selection (Supplementary Fig. 12a) we were able to isolate two amiR-*mail2* lines (amiR-*mail2_4* and amiR-*mail2_5*) that developed further. Both homozygous lines showed around 2.5-fold reduction in *MAIL2* mRNA levels (Supplementary Fig. 12b) and similar developmental phenotypes, including shorter stature, curly leaves, short stems and hypofertility, which were partially suppressed by the introduction of *ring1a/1b* mutations (Fig. 4a and Supplementary Fig. 12c). We confirmed that these phenotypes were caused by *MAIL2* downregulation by restoring MAIL2 function in amiR-*mail2_4* using an amiRNA-insensitive *MAIL2* transgene (Supplementary Fig. 9b, 12d). We evaluated the impact of *MAIL2* downregulation on transcription by performing RNA-seq of amiR-*mail2_4* and amiR-*mail2_5* lines. The transcriptomes of the two lines were highly similar, with differentiated expressed loci almost exclusively comprising genes, either up-(197) or down-regulated (96 genes) (Supplementary Table 2). Downregulated genes were significantly enriched for MAIL2-associated genes and were predominantly found in the top five expression quantiles in WT plants (Fig. 4c,e). These genes showed no enrichment for specific biological pathway and were distinct from those downregulated in *mail1* mutants (Supplementary Fig. 13a). Introduction of amiR-*mail2* into the *mail1* background drastically worsened the *mail1* plant phenotype (Supplementary Fig. 13b), indicating that MAIL1 and MAIL2 function in parallel pathways regulating distinct gene sets. In amiR-*mail2*, H3K27me3 distribution was also affected, and a large fraction of protein-coding genes (75 out of 102) gaining this mark were also bound by MAIL2 (Supplementary Fig. 13c). We further found that genes showing MAIL2 binding to their promoter and downregulation in amiR-*mail2* were associated with TRB1/2/3 in WT plants (Fig. 4f and Supplementary Fig. 13d,e), and specifically gained H3K27me3 upon *MAIL2* downregulation (Fig. 4e,f). We conclude from these results that MAIL2 and MAIL1 are involved in similar pathways antagonizing PcG-silencing to promote expression of largely distinct sets of genes

### M1M and M2M are required to promote expression and oppose H3K27me3 deposition

To assess the functional relevance of M1M and M2M motifs, we employed two complementary approaches based on transgenes. First, we used a transgenic construct in which the promoter of the *SEP3* gene drives *GUS* expression^29^. The *SEP3* promoter contains a telobox that allows PRC2-mediated H3K27me3 deposition through TRB1 binding, resulting in low *GUS* expression^29^. Inserting either M1M or M2M in the *SEP3* promoter between the telobox and the transgene’s TSS significantly enhanced *GUS* expression and reduced H3K27me3 levels (Fig. 5a). In a complementary approach, we generated transgenic lines using the promoters of two endogenous genes: *KPI-1*, a MAIL1/MAIN target, or *HD2A,* a MAIL2 target, both of which are transcriptionally downregulated and enriched for H3K27me3 in their respective mutant backgrounds (Fig. 5b). These promoters naturally contain teloboxes and either M1M (KPI-1 promoter) or M2M (HD2A promoter) motifs. We independently transformed WT plants with constructs carrying either the native promoter or mutated version of M1M or M2M at the most conserved positions. We found that mutating either M1M or M2M motif led to significant downregulation of *GUS* expression and increased H3K27me3 deposition (Fig. 5c,d). These results demonstrate that the integrity of M1M and M2M sequences is required to promote expression and antagonize H3K27me3-associated repression.

**Fig. 5.**
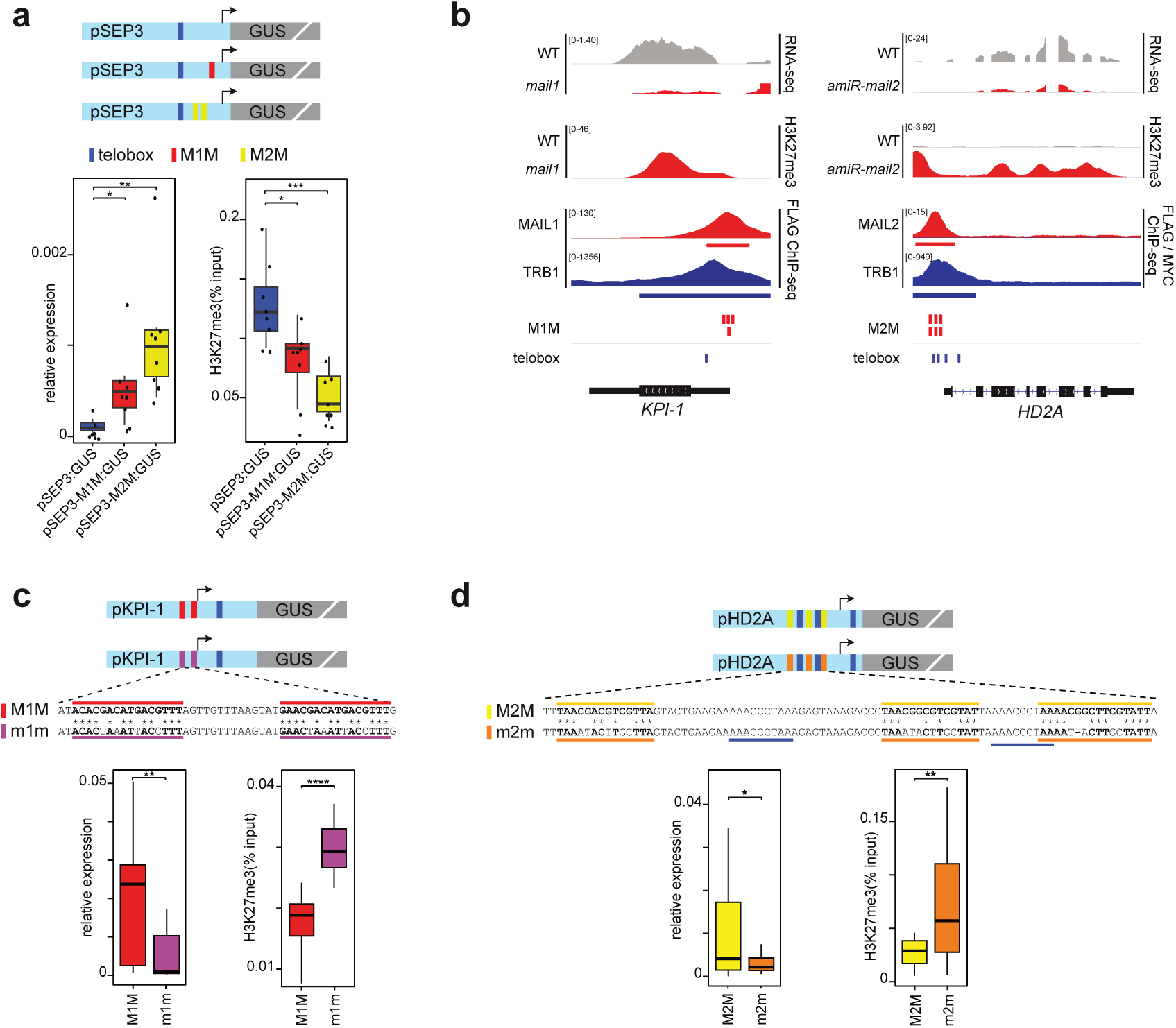
Promoter-associated M1M and M2M are functionally relevant to safeguard genes from PcG-mediated silencing. (a) Schematic representation of the *proSEP3:GUS* original transgene and derivatives following insertion of one M1 or two M2 motifs. Box plots showing relative GUS expression and H3K27me3 accumulation associated to different transgene combinations. For each transgene, data represent two technical replicates on 4 independent transformed lines. Quantitative RT-PCR analysis of *GUS* expression was normalized to *ACT2*. H3K27me3 ChIP-qPCR is reported as % input. Asterisks mark statistically significant differences from *pSEP3:GUS* transgene (unpaired, two-sided Student’s t-test, * *P* < 0.05; ** *P* < 0.01; *** *P* < 0.001; **** *P* < 0.0001). (**b**) Genome browser views of RNA-seq (CPM) and H3K27me3 ChIP-seq (RPGC) profiles at *KPl-1* and *HD2A* loci in the indicated genotypes. MAIL1 and MAIL2-associated patterns are shown together with TRB1-FLAG data (GEO-GSM6895939 retrieved from^30^) and predicted M1M, M2M and telobox motifs. (**c**) Schematic representation of *pPKL1:GUS* transgenes. Most conserved M1M motifs were mutated (m1m) and sequence modifications are shown under the scheme. Telobox motif is also reported in blue. Relative GUS expression and H3K27me3 accumulation were performed as described in (a) except that nine independent lines were analysed for each transgene. (**d**) Schematic representation of *pHD2A:GUS* transgenes showing localization of intact (M2M) and mutated (m2m) motifs. Telobox motifs are indicated in blue and sequence modifications are reported under the scheme. Molecular analyses were performed as described in (a) except that nine independent lines were analysed for each transgene. In box plots, center lines show the medians; box limits indicate the 25th and 75th percentiles; whiskers extend 1.5 times the interquartile range from the 25th and 75th percentiles.

## Discussion

We previously reported that MAIL1, MAIN, and PP7L form a regulatory complex that primarily controls gene expression in Arabidopsis^12–15^, yet the molecular mechanisms remained unclear. Here, we show that mutations in NDX or in the PRC1 components RING1A and RING1B alleviate the developmental and molecular defects associated with MAIL1 depletion, revealing an unexpected functional antagonism between the MAIL1/MAIN pathway and Polycomb silencing.

NDX, a homeodomain-containing protein, interacts with RING1A/B and stabilizes transcriptional states at euchromatic genes^17,19,35^. It was also recently implicated in influencing siRNA accumulation and DNA methylation, predominantly at pericentromeric TEs^31^. While altered DNA methylation can lead to ectopic H3K27me3 deposition at TEs ^36,37^, we did not detect global changes in H3K27me3 patterns in *ndx* mutants (Supplementary Fig. 4). We observed reciprocal co-enrichment of NDX and RING1A/B at their binding sites (Supplementary Fig. 8e,f), consistent with their demonstrated physical interaction^17,19^. Notably, NDX enrichment was detected at protein-coding genes rather than TEs (Supplementary Fig. 8d), contrasting with prior reports of its localization at centromeric and pericentromeric regions^31^. Moreover, NDX and RING1A are co-enriched at MAIL1/MAIN and MAIL2-binding sites (Supplementary Fig. 8e), suggesting that these loci are poised to recruit PRC2. We propose a model whereby MAIL1/MAIN and MAIL2 maintain gene expression by limiting PcG activity at these loci. Upon depletion of MAIL1 or MAIL2, RING1A and NDX may promote PRC2 recruitment, leading to H3K27me3 deposition and gene repression. Additional cofactors such as TRB proteins, known PRC2 recruiters, may further contribute to this process at specific targets.

Interestingly, although VAL1 also recruits PRC1 and PRC2 and cooperates with NDX at *FLC*^19–21^, *val1* mutations do not suppress *mail1* defects but instead exacerbate them (Supplementary Fig. 1e). This suggests that different PRC1/2 recruitment pathways do not contribute equally to MAIL1/MAIN-dependent gene regulation. The aggravated phenotype of *mail1 val1* double mutants may reflect misregulation of distinct gene subsets affected in each background (Supplementary Fig. 14a), although genome-wide binding of VAL1^21^ includes many MAIL1/MAIN targets (Supplementary Fig. 14b), indicating possible context-dependent cooperation.

Our findings establish MAIL1/MAIN and MAIL2 as key regulators that counteract Polycomb-mediated repression to safeguard gene expression. Despite targeting distinct genes via specific promoter motifs (M1M and M2M), these PMD-C proteins likely operate through a shared molecular mechanism. Structural modeling and DNA-binding predictions suggest that the PMD-C domains of MAIL1, MAIN and MAIL2 share a conserved concave surface, a structural feature commonly associated with DNA-binding domains^38^, that includes residues predicted to mediate DNA contact (Supplementary Fig. 15). Moreover, the MAIL1 PMD-C domain shows structural similarity to the DUF1985 domain of the transposon-encoded anti-silencing protein VANC^39^, a known DNA-binding factor^40–42^. This raises the possibility that PMD-C proteins may directly bind DNA, although this interaction remains to be experimentally demonstrated. Alternatively, additional DNA- or chromatin-binding factors may be required to guide or stabilize the interaction of PMD-C with DNA.

We previously linked TE reactivation in *mail1* and *main* mutants to reduced expression of the silencing effector MORC1^15^. Here, we show that *MORC1* is among the genes protected from PcG repression by MAIL1/MAIN. In *mail1 ndx* and *mail1 ring1a/b* mutants, TEs are largely re-silenced, likely due to restored expression of MORC1 along with upregulation of its paralog MORC2 (Fig. 2b and Supplementary Fig. 3). Many genes are upregulated in *mail1*, possibly due to the downregulation of transcriptional repressors. However, we did not identify any clear candidate. *MORC2* is unlikely to be involved, as its expression remains very low in both *mail1* and WT (Supplementary Fig. 3a), and *morc2* mutants show no gene deregulation^24^. *MORC1* is also most likely not responsible, as the genes upregulated in *mail1* and *morc1* are largely distinct (Supplementary Fig. 14c), and *MORC1* overexpression in *mail1* has minimal impact^15^. Alternatively, gene upregulation in *mail1* may result from changes in higher-order chromatin architecture associated with loss of MAIL1 function^12^.

*MAIL1* and *MAIN* likely evolved from a duplication of *MAIL2* that occurred specifically within the Brassicaceae lineage^13^. The sequence similarity between the M1M and M2M motifs suggests that the palindromic M2M motif may have evolved into M1M at the same time. *MAIL2* appears conserved at least in all eudicot plants^13^, raising the possibility that this PMD protein-DNA motif regulatory module may be conserved across a broad range of plant species. Only a subset (about 20-40%) of M1M and M2M motifs are bound by MAIN/MAIL1 or MAIL2 (Supplementary Fig. 13f), suggesting that additional regulatory layers restrict their binding to specific active genes. Future studies will address this point.

We previously defined four clades of PMD proteins (A1, A2, B and C). PMD-As are encoded by retrotransposons, while PMD-B are found in protein-coding genes. MAIL1, MAIN and MAIL2 belong to the PMD-C clade, which comprises protein synthesized from *bona fide* host genes as well as DNA transposon-encoded proteins from various monocots and eudicots plant species^12^ (Supplementary Fig. 16 and Supplementary doc. 1). The predicted structures of MAIL1, MAIN and MAIL2 PMD-C domains are highly similar (Supplementary Fig. 10) and, as mentioned above, share interesting structural homology with the DUF1985 domain of the DNA transposon-encoded VANC proteins^39^, which function as an anti-silencing factor antagonizing DNA methylation ^40–42^. Recent studies indicate that H3K27me3 plays a role in TE repression in a broad range of eukaryotes, including Arabidopsis, suggesting an ancestral role of Polycomb silencing in controlling TE activity^43,44^. This raises the intriguing possibility that TE-encoded PMD-C proteins also function as Polycomb anti-silencing factors, a hypothesis we are currently testing. Such a connection between transposon biology and Polycomb silencing regulation could provide valuable insights into the coevolution of genome defense and gene expression control mechanisms.

## Methods

### Plant Material

Mutants *mail1-1* (GK-840E05)^27^, *main-2* (GK-728H05)^45^*, mail2-1* (SAIL_1153_F07), *ndx-4* (WiscDsLox344A04)^35^*, ring1a-2* (SAIL_393_E05)*/1b-3* (SAIL_520_B04)^18^*, clf-29 (*SALK_021003)^46^, *swn-7 (*SALK_109121)^6^ and *val1-2* (SALK_088606C)^21^ are all in the Col-0 background. The transgenic L5 line^16^ was kindly provided by H. Vaucheret (IJPB, France).

Design of the amiRNA (amiR) construct directed against *MAIL2* transcript was done using the WMD3-Web MicroRNA Designer (http://wmd3.weigelworld.org) and the amiR-*MAIL2* fragment cloned into the BIN-HYG-TX plasmid under the control of p35S promoter.

The *pMAIL1:gMAIL1-FLAG*, *pMAIL1:gMAIL1-MYC*, *pMAIN:gMAIN-FLAG*, *pMAIN:gMAIN-MYC* and *SEP3pro:GUS* constructs were described previously^13,29^. For the *pMAIL2:gMAIL2-9xMYC* construct, two genomic fragments encompassing the *MAIL2* gene (−1834 to +2448 relative to ATG codon; A designated as +1) were PCR-amplified from Col-0 genomic DNA. PCR primers, with several point mutations to *MAIL2* sequence, allowed to reconstitute a *MAIL2* transgene bearing silent mutations and insensitive to amiR-*MAIL2* targeted degradation (Supplementary Fig. 9b); recombination-based technology (NEB, NEBuilder HiFi DNA Assembly Master Mix #2621) was then used to replace *MAIL1*-specific sequences of *pMAIL1:gMAIL1-MYC* vector by the *MAIL2* transgene fragment. The *pNDX:gNDX* fragment was PCR amplified from Col-0 genomic DNA and region from −1631 to +5177 relative to ATG codon was cloned into the PMDC123 vector. This *pNDX:gNDX* construct was used to produce a Nter-FLAG version of NDX by inserting three repeats of the FLAG tag in frame with the second amino acids of the NDX coding sequence. A Cter-9xMYC tag version was obtained through a recombination-based approach similar to the one described for generating the *pMAIL2:gMAIL2-9xMYC*. For *KPI-1:GUS* and *HD2A:GUS* transgenes, proximal gene promoter and 5’UTR amplicons of 459bp and 357bp respectively were fused to *GUS* coding sequence and cloned into the PMDC123 vector. M1M and M2M were mutated using *pKPI-1:GUS* or *pHD2A:GUS* as matrix and sequence modifications are described on Fig. 5. Insertion of M1M (−321 to −306 relative to ATG) or M2M (−460 to −423 relative to ATG) in *proSEP3:GUS* was done by inverse PCR.

All transgenic constructs were introduced in plants through agroinfiltration by floral dipping^47^ and T1 transformants were isolated following adequate antibiotic selection.

Plants were grown on soil (60% relative humidity) or *in vitro* under long-day conditions (16 h light, 8 h dark) at 23 °C.

### Mutagenesis and mapping

Ethyl methanesulfonate (EMS) mutagenesis of the L5/*mail1-1* mutant line was done as previously described^12^. To isolate suppressors of the *mail1* mutation, a double screen was performed based on: 1/ visual amelioration of plant phenotype 2/ GUS staining to confirm L5 transgene resilencing; GUS assay was done by dissecting one leaf per M2 plant candidate and leaves were incubated at 37 °C for 24 h in 400µg ml^-1^ 5-bromo-4-chloro-3-indolyl-b-D-glucuronic acid, 10 mM EDTA, 50 mM sodium phosphate buffer pH 7.2, 0.2% triton X-100. The *ndx-7* mutant allele was identified following mapping-by-sequencing using genomic DNA from a pool of 47 individual F2 mutant plants segregating from a cross with Ler

### RNA isolation and analyses

Total RNA was extracted from aerial tissues of 12-day-old plants or immature flower buds using RNAzol RT (RN190, Molecular Research Center, Inc). For real time RT-qPCR, 0.5 µg of total RNA was DNase treated and reverse transcribed using the PrimeScript RT reagent kit with gDNA eraser (Perfect real time) (TaKaRa) in a final volume of 20 µl and one microlitre of cDNA was used for subsequent amplification using SYBR Premix Ex Taq II (Tli RnaseH Plus) (TaKaRa) and Real-Time LightCycler 480 system (Roche) in a final reaction volume of 15 µl. Data were normalized to a reference gene and analyzed according to the 2^−ΔΔCt^ method. Means and standard errors of the mean were calculated from independent biological samples. Primers used are listed in Supplementary Table 3.

For RNA-seq, DNase and purification were performed using RNA Clean & Concentrator-5 kit (Zymo Research). Libraries were generated from about one µg total RNA and sequenced as PE100 or PE150 reads on a DNBSEQ platform at Beijing Genomics Institute (Hong Kong).

### ChIP-seq

About two grams of aerial two-week-old rosettes were collected and incubated 15 min under vacuum in 25 ml of freshly prepared cross-linking buffer (10 mM phosphate buffer pH 7, 100 mM NaCl, 0.1 M sucrose, 1% formaldehyde). 2 M glycine was added to stop cross-link reaction (0.125 M final) and incubated an additional 5 min under vacuum. Tissues were then washed three times with 10 mM phosphate buffer pH 7, 100 mM NaCl, 0.1 M sucrose, pooled in 50 ml falcon tube and snap frozen in liquid nitrogen. Cross-linked tissues were ground into fine powder by adding 7 ceramic balls to falcon tube and vortexing, and resuspended into 25 ml of extraction buffer 1 (0.4 M Sucrose, 10 mM Tris-HCl pH 8, 10 mM MgCl2, 5 mM β-mercaptoethanol, 0.1 mM PMSF, 1X complete EDTA-free cocktail inhibitor (Roche)) at 4°C. Alternatively, collected tissues were ground into fine powder with liquid nitrogen and samples were resuspended with 25 ml Hepes buffer containing 1% formaldehyde (50 mM Hepes pH 8, 1 M sucrose, 5 mM KCl, 5 mM MgCl2, 0.6% triton X-100, 0.4 mM PMSF, 1% formaldehyde, 1X complete EDTA-free cocktail inhibitor), and incubated for 10 min at RT. Crosslinking was stopped with 2 M glycine (0.125 M final), inverting several times. The lysates were filtered through one layer of Miracloth and the nuclei were collected by centrifuge at 4°C with 4000 rpm for 20 min. The nuclei were resuspended with extraction buffer 2 (0.25 M Sucrose, 10 mM Tris-HCl pH 8, 10 mM MgCl_2_, 1% triton X-100, 5 mM β-mercaptoethanol, 0.1 mM PMSF, 1X complete EDTA-free cocktail inhibitor), centrifuge with 14000 g at 4°C for 10 min, and then resuspended with 400 µl extraction buffer 3 (1.7 M Sucrose, 10 mM Tris-HCl pH 8, 0.15% Triton X-100, 2 mM MgCl_2_, 5 mM β-mercaptoethanol, 0.1 mM PMSF, 1X complete EDTA-free cocktail inhibitor). The nuclei suspension was loaded over a cushion of 400 µl extraction buffer 3 and microtubes were centrifugated with 14000 g for 1 h at 4°C. For chromatin shearing using Bioruptor UCD-200 (Diagenode), nuclei were resuspended in 250 µl of lysis buffer (50 mM Tris-HCl pH 8, 10 mM EDTA pH 8, 1% SDS, 1X complete EDTA-free cocktail inhibitor) and 25 cycles of 25 s ‘ON’, 1 min 15 s ‘OFF were applied at 4°C. Following centrifugation, chromatin was split into two fractions of 80 µl and 9 vol. of ChIP dilution buffer added (1.1% Triton X-100, 16.7 mM Tris-HCl pH 8, 1.2 mM EDTA pH 8, 167 mM NaCl, 1X complete EDTA-free cocktail inhibitor) before proceeding to preclearing and IP as described hereafter. For Covaris fragmentation, nuclei were resuspended with 1 ml of covaris buffer (20 mM Tris pH 8, 2 mM EDTA, 0.1% SDS, 1X complete EDTA-free cocktail inhibitor) for 30 min at 4°C on a turning wheel. The lysate was transferred into Covaris milliTube Glass 1 ml AFA Fiber and chromatin was mainly sheared to 200-400 bp fragments using S220 Focused-ultrasonicator with the following conditions: for histone marks analysis, PIP175, CBP200, DF20, T 5°C, 5 min; for MAIL2-MYCtag, PIP140, CBP200, DF10, T 5°C, 6 min. Sheared chromatin was centrifuged with 21000 g for 10 min at 4°C and the supernatant was split into two fractions of 440µl. After adding 0.5 volume of covaris dilution buffer (20 mM Tris-HCL pH 8, 2 mM EDTA, 0.1% SDS, 450 mM NaCl, 3% Triton X-100, 1X complete EDTA-free cocktail inhibitor), chromatin was precleared with 30 µl of magnetic Protein A and Protein G Dynabeads (Invitrogen) for 1 h at 4°C on a turning wheel. Twenty µl was saved as input and the remaining supernatant was then incubated at 4°C overnight with specific antibodies (5 µg of H3K27me3 Ab, Diagenode C15410069 or Millipore 07-449; 10 µl H2AUb Ab, Cell Signaling Technology ref.8240; 10 µl Myc-Tag Ab, Cell Signaling Technology ref.2276 or ref.2278; 3.5 µl monoclonal ANTI-FLAG M2 Ab, Sigma F1804-1MG). Next, 40 µl of magnetic Protein A and Protein G Dynabeads were added and the mixture was rotated at 4°C for 3-4h. The beads were washed for 5 min rotating at 4°C with low salt solution twice (0.1% SDS; 1% Triton X-100, 2 mM EDTA pH 8, 20 mM Tris-HCl pH 8, 150 mM NaCl), high salt solution (0.1% SDS; 1% Triton X-100, 2 mM EDTA pH 8, 20 mM Tris-HCl pH8, 500 mM NaCl), LiCl solution (0.25 M LiCl, 1% IGEPAL CA630, 1% deoxycholic acid, 1 mM EDTA, 10 mM Tris-HCl pH8) and TE solution (10 mM Tris-HCl pH 8, 1 mM EDTA pH 8). A second TE wash was performed after transferring beads to a new DNA low-bind Eppendorf microtube. The chromatin was eluted with 250 µl elution buffer twice (1% SDS, 0.1 M NaHCO3) at 65°C for 15 min on a thermomixer with shaking (950 rpm) and subjected to reverse crosslinking by adding 51 µl of reverse crosslink mix (2 M NaCl, 0.1 M EDTA pH 8, 0.4 M Tris-HCl pH 8, 0.4 mg/ml proteinaseK) for 3 h at 50°C then overnight at 65°C. Following digestion with 10 µg RNaseA for 30 min at RT°, DNA was purified using ChIP DNA clean and concentrator kit (Zymo ZD5205). For MAIL2-MYC, five independent IP experiments (equivalent 10 g tissues) were pooled.

For immature flower buds, three tubes of about 0.15 g material were harvested and snap-frozen in liquid nitrogen. Following tissue disruption using Qiagen TissueLyser, samples were resuspended in 1.8 ml Hepes Buffer containing 1% formaldehyde, pooled and further processed as described above with covaris fragmentation at 5°C, PIP140, CBP200, DF10 for 6 min. IP was then performed using 4 µg of antibody to H3K27me3 (Diagenode, C15410069). Libraries were generated and sequenced as PE100 reads on a DNBSEQ platform at Beijing Genomics Institute (Hong Kong).

### ChIP-qPCR

Three tubes of about 0.15 g of aerial 12-day-old rosettes were snap frozen in liquid nitrogen and processed as described above for floral buds, except that nuclei were resuspended in 130 µl covaris buffer and sonicated using microtube-130 AFA Fiber. Each IP was performed using 2.1 µg of H3K27me3 antibody. (Diagenode, C15410069) and one of 15 µl purified DNA was used for real-time PCR amplification using SYBR Premix Ex Taq II (Tli RnaseH Plus) (TaKaRa) and Real-Time LightCycler 480 system (Roche) in a final reaction volume of 15 µl. Primers used are listed in Supplementary Table 3.

### CUT&Tag

About 0.15g of WT, *ndx-7*::gNDX-9xMYC or *ndx-7*::g3xFLAG-NDX immature flower buds was snap frozen in liquid nitrogen, grounded to a fine powder using Qiagen TissueLyser followed by a light *in vitro* cross-linking for 2 min at RT with 1.8ml ChIP extraction buffer 1 containing 0.2% formaldehyde. Cross-linking was stopped with 122.5µl of 2M glycine rotating for 10 min at RT. Lysate was filtrated through two layers of Miracloth and centrifugated for 20 min at 3000 g and 4°C. Nuclei were resuspended into 1ml of ChIP extraction buffer 2 and centrifugated with 12000 g at 4°C for 10 min; this step was repeated for a total of 3 washes until the nuclei pellet appears totally white. The resulting pellet was resuspended in 250 µl ChIP extraction buffer 2, then loaded over a cushion of 250 µl ChIP extraction buffer 3 and centrifugated with 16000 g for 1 h at 4°C. Nuclei were resuspended in 1 ml Nuclear Extraction (NE) buffer (20 mM Hepes pH 7.9, 10 mM KCl, 0.1% triton X-100, 20% glycerol, 0.5 mM Spermidine, 1X complete EDTA-free cocktail inhibitor), incubated for 10 min on ice followed with 3 min centrifugation for 600 g at 4°C. The extract was resuspended in 500 µl NE buffer and centrifuged 3 min for 600 g at 4°C. The nuclei fraction was finally resuspended in 500 µl NE buffer and nuclei concentration was estimated by counting using Neubauer chamber. Each CUT&Tag reaction was then performed on 400 000 nuclei essentially as described in the Epicypher CUTANA CUT&Tag protocol (v1.7) and using 0.5µl monoclonal Myc-Tag Ab (Cell Signaling Technology ref.2276) or 0.5 µl monoclonal ANTI-FLAG M2 Ab (Sigma F1804-1MG). Libraries were amplified with 18 PCR cycles and > 75 bp DNA fragments were cleaned through two rounds of purification using 1.3x, then 0.8x AMPure Beads (Beckman Coulter A63880). Libraries were pooled and sequenced as PE100 reads on a DNBSEQ platform at Beijing Genomics Institute (Poland).

### RNA-seq analysis

Reads were trimmed and filtered for quality and adapter contamination using Trim Galore (v.0.6.7, Babraham Institute) and aligned to the *Arabidopsis* reference genome (TAIR10) genome using STAR (v.2.7.9a)^39^. Reads aligning equally well to more than one position in the reference genome were discarded, and probable PCR duplicates were removed using MarkDuplicates from the Picard Tools suite^49^. Sample tracks were generated using deeptools (v.3.5.1)^50^ bamCoverage with the option normalizeUsing CPM and binSize 1. Subsequent read counting for each gene and TE was performed using featureCounts (v.2.0.3)^51^ using the Araport11 genome annotation. Annotated TEs overlapping strongly (> 80%) with an annotated TE gene were considered TE genes, and the TE annotation was discarded. Differential expression analysis was performed using DESeq2 (v.1.36.0)^52^ with absolute fold-change ≥ 4 and Benjamini-Hochberg adjusted *p*-value < 0.05 were considered differentially expressed. For DEseq2 analysis of amiR-*mail2* hypomorphic lines, criteria of absolute fold change ≥2 and Benjamini-Hochberg adjusted *p*-value < 0.05 were retained. All plots were generated using ggplot2 (v.3.4.4).

### ChIP-seq analysis

Reads were trimmed and filtered using Trim Galore (v.0.6.7, Babraham Institute) or Trimmomatic (v.0.39)^53^, and aligned to the TAIR10 *Arabidopsis* reference genome using bowtie2 (v.2.4.2)^54^, allowing only uniquely mapped reads with perfect matches. PCR duplicates were discarded using MarkDuplicates from the Picard Tools suite or markdup from Sambamba (v.0.8.2)^55^. Sample tracks were generated using deeptools (v.3.5.1)^50^ bamCoverage with the option normalizeUsing RPGC and binSize 1. Peaks calling was performed with MACS2 (v.2.2.7.1)^56^ using corresponding inputs as a background and -- nomodel and –extsize 200 options or MACS3 (v.3.0.0b1)^56^ using --model and –call-summits, q-value < 0.01, and peaks called in the anti-MYC or anti-FLAG WT controls were removed. MAIL1/MAIN bound regions are designed as the summit of peaks plus 100bp on each side.

### CUT&Tag data processing

Raw paired-end reads were trimmed using Trim Galore (v.0.6.7) with default parameters. Trimmed reads were aligned to the TAIR10 reference genome using Bowtie2 (v.2.4.2 in --very-sensitive-local mode, with insert size constraints (−I 10 -X 700), and with --no-mixed and --no-discordant options to restrict alignments to properly paired reads. Reads mapping to organellar genomes (chloroplast or mitochondria) were filtered out, and only uniquely mapping reads with a mapping quality ≥ 10 were retained. BAM files were then sorted and indexed and insert size distributions were computed using Picard’s *CollectInsertSizeMetrics*. For sample normalization, exogenous *E. coli* DNA carried by the pA-Tn5 transposase was used as a spike-in control. Clean reads were aligned to the *E. coli* genome using Bowtie2, and a scaling factor was computed for each sample as the ratio between the number of *E. coli* reads in the sample and the smallest *E. coli* read count observed among all samples. These scaling factors were used to normalize read coverage when generating signal tracks (bigWig format) using *bamCoverage* of deepTools (v.3.5.1), with a bin size of 5 bp.

### Motif discovery

DNA sequences of MAIL1/MAIN and MAIL2 ChIP-seq peaks and of gene proximal promoter regions (+/- 500 pb surrounding TSS) were extracted using bedtools (v.2.31.1) getfasta. Conserved motifs were identified using STREME v5.5.5^57^ and the shuffled input sequences option with the following parameters: streme --verbosity 1 --oc streme_out -dna --minw 6 --maxw 16 --order 2 --bfile ./background --seed 0 --align center --time 4740 -- totallength 4000000 --evalue --thresh 0.05, or using MEME v5.5.5^58^ with the Classic mode option and the following parameters: meme -oc meme_out -objfun classic -mod anr -csites 10000 -nmotifs 10 -minw 6 -maxw 15 -markov_order 0 -dna -revcomp -evt 0.05. Differentially enriched DNA motifs were recovered using the MEME DE mode option and an equivalent control set; relative enrichment ratio was evaluated with the SEA program of the MEME suite. Genomic distribution of M1M and M2M identified from ChIP peaks was determined using FIMO v5.5.5^59^ with *P* value match < 1e-05 and default parameters. Genomic localization of telobox elements was determined using FIMO v5.5.5^59^ with p-value match < 1e-05 and default parameters and the consensus sequence “AAACCCTAR”.

### Gene ontology analysis

GO term enrichment analysis was performed using ShinyGO 0.82^60^ and the *Arabidopsis thaliana* Araport11 annotation. *P*-value was derived from hypergeometric distribution, adjusted using the FDR (false discovery rate) method. Results with a *P*-value < 1e-05 were considered as significant.

### Protein structure and DNA-binding potential prediction

The structures of the PMD-C domains of MAIL1, MAIN and MAIL2 were modeled using AlphaFold 3 ^61^. Potential DNA-binding residues of MAIN, MAIL1 and MAIL2 were predicted using the geometry-aware binding site predictor GPSite ^62^. For each PMD-C model, prediction scores were mapped onto the corresponding residues for visualization using UCSF ChimeraX v. 1.9 ^63^.

### Phylogenetic analysis

Candidate MuDR-encoded PMD sequences were identified through tBlastN analysis of Repbase database^64^ using the MAIL2 PMD-C region as query (Supplementary doc. 1). The “PhyML+SMS/OneClick” pipeline^65^ (https://ngphylogeny.fr/) with default parameters was then used to build a phylogenetic tree based on MuDR-associated PMD and including Arabidopsis gene-encoded PMD-B and -C sequences. Resulting tree was exported using the online Interactive Tree Of Life (ITOL) tool. PMD sequences used to build the tree are reported in Supplementary Table 4.

## Data availability

The data supporting the findings of this study are available within the article and its Supplementary Information. High throughput sequencing data has been deposited in the Gene Expression Omnibus (GEO) database and can be accessed with the accession number GSE278560 (accessible using the token *ktghyqoafluttcz* during the review process). All data are available from the corresponding author upon reasonable request.

## Acknowledgments

We thank F. Turck (Max Planck Institute for Plant Breeding Research) for kindly providing the *SEP3pro:GUS* construct. We are also grateful to S. Marquardt (Department of Plant and Environmental Sciences, University of Copenhagen) for his help in setting up the CUT&Tag assay. Work in the Mathieu laboratory was supported by CNRS, Inserm, Université Clermont Auvergne core funding, a grant from the iSITE CAP2025 (to M.O.) and grants from the Agence Nationale de la Recherche (ANR-20-CE12-0009 & ANR-23-CE20-0012 to O.M.). Work in the Moissiard laboratory was supported by CNRS and University of Perpignan Via Domitia (UPVD) core funding, and by grants from the Agence Nationale de la Recherche (ANR-23-CE20-0012 to G.M). This study was also supported by “Laboratoires d’Excellences” (LAbEX) AGRO 2011-LABX-002 (under I-Site Muse framework) coordinated by Agropolis Foundation (ID 2101-009 to G.M), region Occitanie PhD grant to L.J, and by the “Laboratoires d’Excellences (LABEX)” TULIP (ANR-10-LABX-0041)” and “École Universitaire de Recherche (EUR)” TULIP-GS (ANR-18-EURE-0019). The funders had no role in study design, data collection and analysis, decision to publish, or preparation of the manuscript.

## Author contributions

T.P., L.J., G.M. and O.M. conceived the study. T.P., L.J., M.O., G.D., M.-N.P.-P., C.C., J.D., N.P., G.M. and O.M. conducted laboratory experiments. T.P., L.J., G.M. and O.M interpreted the data. T.P. and O.M. drafted the manuscript. T.P., L.J., G.M. and O.M. edited the manuscript. G.M and O.M. coordinated the research. The authors read and approved the final manuscript.

---

**Supplementary Table 1. List of H3K27me3-enriched genes (Log2 FC > 1; *P*-value < 0.01) in *mail1* compared to WT background.**

**Supplementary Table 2. Lists of differentially expressed loci in amiR-*mail2*_4 and _5 mutant lines.**

**Supplementary Table 3. List of qPCR primers used in this study.**

**Supplementary Table 4. List of PMD sequences used for phylogenetic analyses.**

**Supplementary Table 5. DNA-binding prediction scores for amino acid residues of MAIN, MAIL1 and MAIL2.**

**Supplementary doc. 1. Best results of TBLASTN search on Repbase**^64^ **using MAIL2 PMD-C domain as query.** Only results with e-values < 1e-10 were reported. Dicot species: ArHy *Arachis hypogaea*, AvMa *Avicennia marina*, Cba *Capsicum baccatum*, Daca *Daucus carota*, Gar *Gossypium arboretum*, GR *Gossypium raimondii*, GM *Glycine max*, Ipa *Ilex paraguariensis*, LU *Linum usitatissimum*, MT *Medicago truncatula*, NS *Nicotiana sylvestris*, SeTo *Senna tora*, STu *Solanum tuberosum*, VV *Vitis Vinifera*. Monocot species: DRot *Dioscorea cayennensis subsp. Rotundata*, OS *oryza Sativa*, SBi *Sorghum bicolor*, SHS *Saccharum hybrid cultivar*, Sit *Setaria italica*, Tae *Triticum aestivum*, ZM *Zea mays*. Basal angiosperm species: ATr *Amborella trichopoda*.

