## Supplementary Figures for "Plant Mobile Domain protein-DNA motif modules counteract Polycomb silencing to stabilize gene expression"

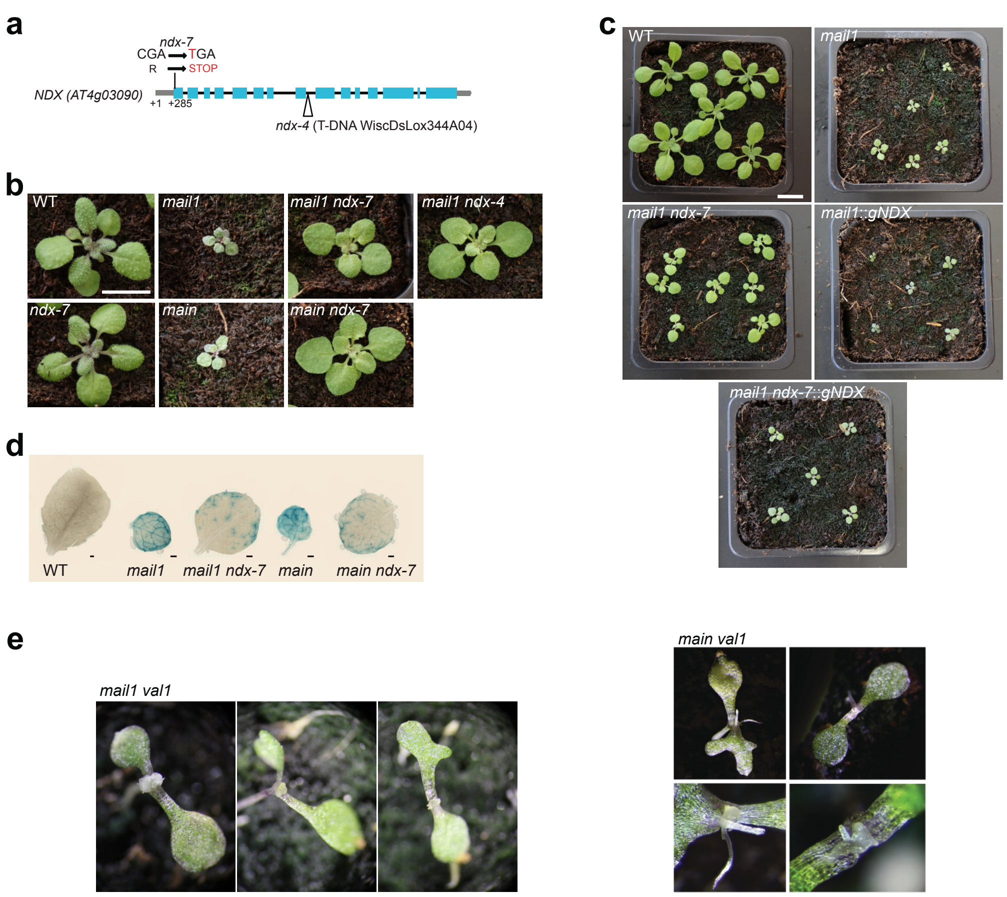


**Supplementary Fig. 1. *mail1*- and *main*-associated phenotypes are suppressed in *ndx* but not in *val1* mutant backgrounds. (a)** Representation of *NDX* describing the insertion (*ndx-4*) and point mutation (*ndx-7*; this study) alleles. C to T mutation for *ndx-7* created an early STOP codon. Position is indicated relative to the transcription start site (+1). (**b**) Developmental phenotype of 3-week-old WT, *mail1*, *main*, *ndx-7*, *mail1 ndx-7*, *mail1 ndx-4* and *main ndx-7* seedlings. Scale bar is 1 cm. (**c**) Photos of two-week-old seedlings of indicated genotype. Scale bar is 1 cm. (**d**) Histochemical staining for GUS activity in leaves from plants of the indicated genotypes. Scale bar is 500 µm. (**e**) Developmental phenotypes observed for *mail1 val1* and *main val1* seedlings showing abnormal differentiated structures emerging from apical meristem.

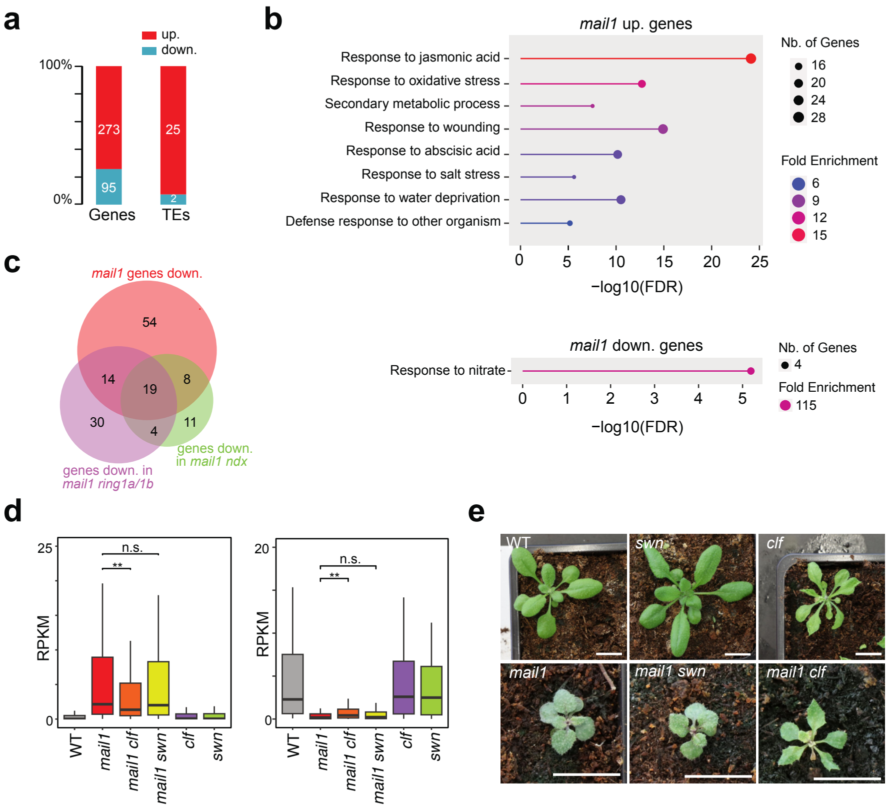


**Supplementary Fig.2. *clf* and *swn* mutations poorly suppress *mail1*- associated molecular and developmental defects. (a)** Fraction and number of TEs and genes significantly up- or down-regulated in *mail1* seedlings versus WT (Log2 FC ≥ 2; Padj < 0.05). (**b**) Results of GO enrichment term analysis for biological processes using the sets of *mail1*-misregulated genes (*P*-value (FDR) < 1e-05). (**c**) Venn diagrams showing overlaps of genes downregulated in *mail1*, *mail1 ndx* and *mail1 ring1a/1b* mutant backgrounds. (**d**) Transcript accumulation (in RPKM) at *mail1*-upregulated (left panel) and *mail1*-downregulated (right panel) genes in the indicated genotypes. Means of two biological replicates were computed. Center lines show the medians; box limits indicate the 25th and 75th percentiles; whiskers extend 1.5 times the interquartile range from the 25th and 75th percentiles. Statistical significances of *mail1 clf* or *mail1 swn* values compared to *mail1* are indicated (Wilcoxon rank sum test with continuity correction; two-sided; n.s. non-significant; ** $P < 0.01$). (**e**) Photos of three-week-old seedlings of indicated genotype. Scale bar is 1 cm.

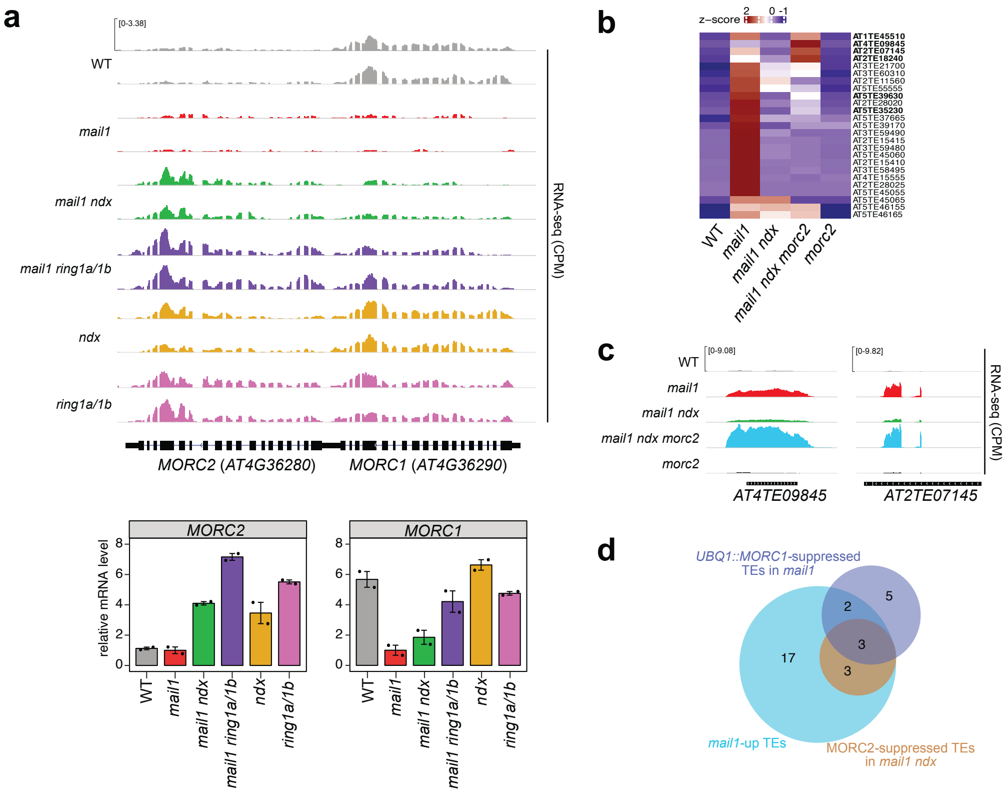


**Supplementary Fig.3. *MORC2* upregulation in *mail1 ndx* plants participates to TE resilencing.** (**a**) Genome browser views of RNA-seq (CPM) profiles at *MORC1* and *MORC2* loci in the indicated genotypes. Two biological replicates are shown. Barplots below show *MORC1* and *MORC2* mean transcript level in the indicated genotypes relative to *mail1* (set to 1). (**b**) Heatmaps of RNA-seq expression z-score computed for *mail1*-upregulated (-up) TEs across the indicated genotypes. Means of two biological replicates were computed. TEs showing significant release of expression in *mail1 ndx morc2* compared to *mail1 ndx* (Log2 FC > 2) are indicated in bold. (**c**) RNA-seq genome browser views at two *mail1*-up TEs in the indicated genotypes. (**d**) Venn diagram analyses showing the overlap between *mail1*-up TEs, *mail1*-up TEs suppressed in *mail1 ndx* due to increased MORC2 expression and *mail1*-up TEs suppressed upon ectopic *UBQ1::MORC1* expression[15].

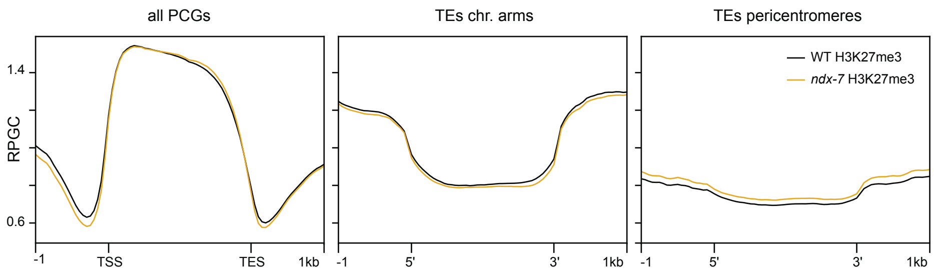


**Supplementary Fig. 4. NDX depletion does not impact overall H3K27me3 distribution.** Metaplots of H3K27me3 accumulation (in RPGC) at protein-coding genes (left) and TEs located in chromosomal arms (middle) or pericentromeric regions (right) in WT and *ndx-7* mutant backgrounds.

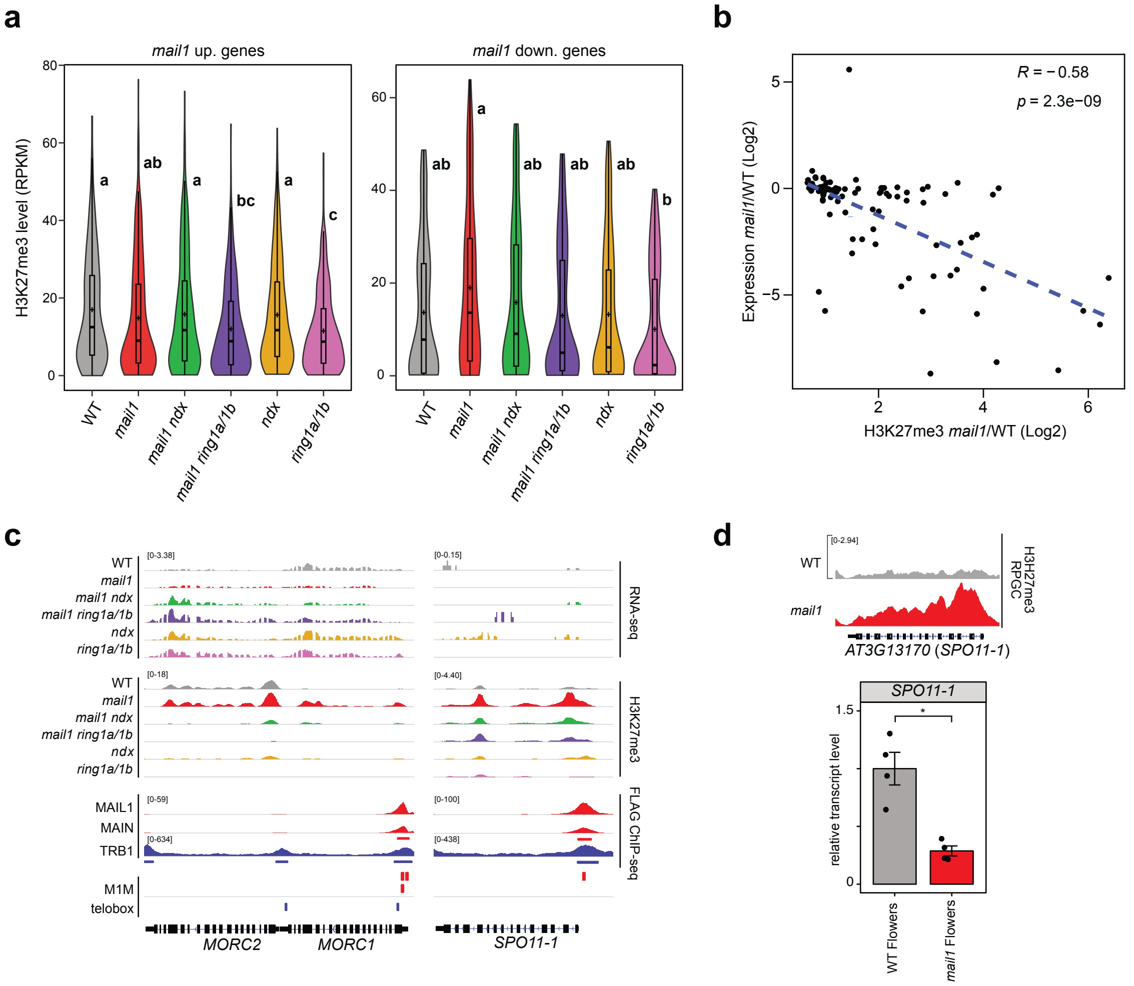


**Supplementary Fig. 5. MAIL1 complexes prevent H3K27me3 incorporation at different loci** (**a**) Violin plot inlaid with boxplot showing H3K27me3 accumulation (in RPKM) at *mail1*-upregulated (up.) and *mail1*-downregulated (down.) genes in the indicated genotype. For box plots, cross and center lines indicate the means and medians, respectively; whiskers extend 1.5 times the interquartile range from the 25th and 75th percentiles. Statistical analysis was performed with analysis of variance (ANOVA); lowercase letters indicate significant differences between groups using Tukey's post hoc tests ($P < 0.05$) (**b**) Scatter plot showing negative correlation between gain of H3K27me3 and gene expression in *mail1*. Log2 fold changes were computed for genes with increased H3K27me3 level in *mail1* vs WT (DESeq2, *P*-value < 0.1) and accumulating RNA in at least one of the genotype at this developmental stage. *R*: Spearman rank correlation coefficient, *p*: significance level. (**c**) Genome browser views of RNA-seq (CPM) and H3K27me3 ChIP-seq (RPGC) profiles at *mail1* H3K27me3-enriched genes in the indicated genotypes. MAIL1-FLAG and MAIN-FLAG associated patterns are shown together with TRB1-FLAG data (GEO-GSM6895939 retrieved from[30]). Predicted Mail1 (M1M) and telobox motifs are indicated. (**d**) Genome browser view of H3K27me3 levels at *SPO11-1* gene in WT and *mail1* immature flowers (top). Transcript levels measured by RT-qPCR (bottom) are normalized to *ACT2* and further normalized to WT; values represent means from four biological replicates ± s.e.m and statistically significant difference is indicated (unpaired, two-sided Student's t-test, * $P < 0.05$).

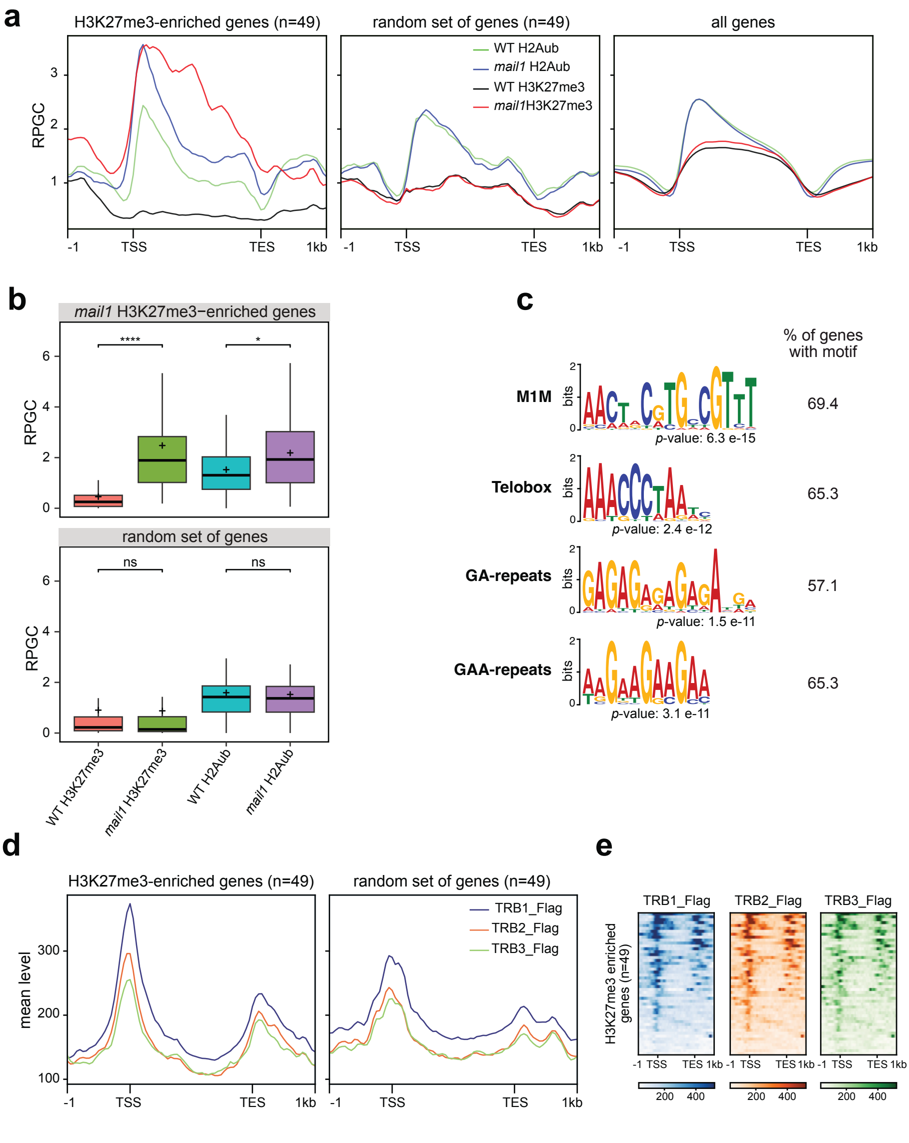


**Supplementary Fig. 6. Ectopic H3K27me3 deposition is detected at TRBs-enriched genes and correlates with increased levels of H2AUb incorporation.** (**a**) Metagene plots of H3K27me3 and H2Aub accumulation (in RPGC) in the indicated genotypes at *mail1* H3K27me3-enriched genes compared to an equivalent random set of genes or to all Arabidopsis genes. (**b**) Box plots illustrating H3K27me3 and H2Aub accumulation (in RPGC) at *mail1* H3K27me3-enriched genes (upper panel) compared to a random set of genes (bottom panel) in the indicated genotypes. Cross and center lines indicate the means and medians, respectively; box limits indicate the 25th and 75th percentiles; whiskers extend 1.5 times the interquartile range from the 25th and 75th percentiles. Statistical significances of *mail1* values compared to WT are indicated (Wilcoxon rank sum test with continuity correction; two-sided; n.s. non-significant; * $P < 0.05$; **** $P < 0.0001$). (**c**) Sequence logos of most conserved DNA motifs detected by STREME analysis within the 1kb region surrounding TSS of H3K27me3-enriched genes in *mail1*. Percentage of regions exhibiting each motif is indicated. (**d**) Metagene plots showing TRB1/2/3 levels (data from[30]) at *mail1* H3K27me3-enriched genes or at an equivalent random set of genes. (**e**) Plot heatmaps illustrating TRB1/2/3 enrichment (data from[30]) at TSS of *mail1* H3K27me3-enriched genes.

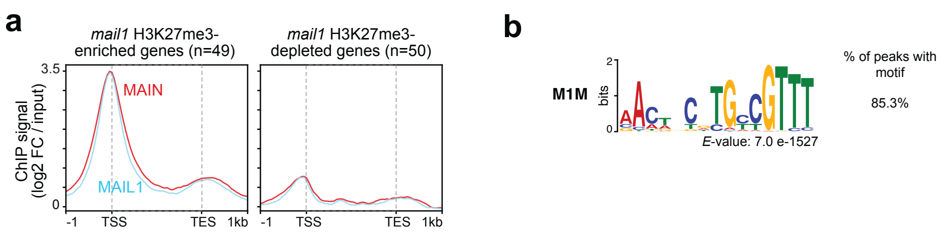


**Supplementary Fig. 7. MAIL1 and MAIN mainly co-localize at M1M-enriched gene TSS.** (**a**) MAIL1 and MAIN signals (LOG2 FC vs input) at *mail1* H3K27me3-enriched or depleted genes. (**b**) Sequence logo of a highly enriched MEME-predicted motif among the 626 MAIL1/MAIN peaks.

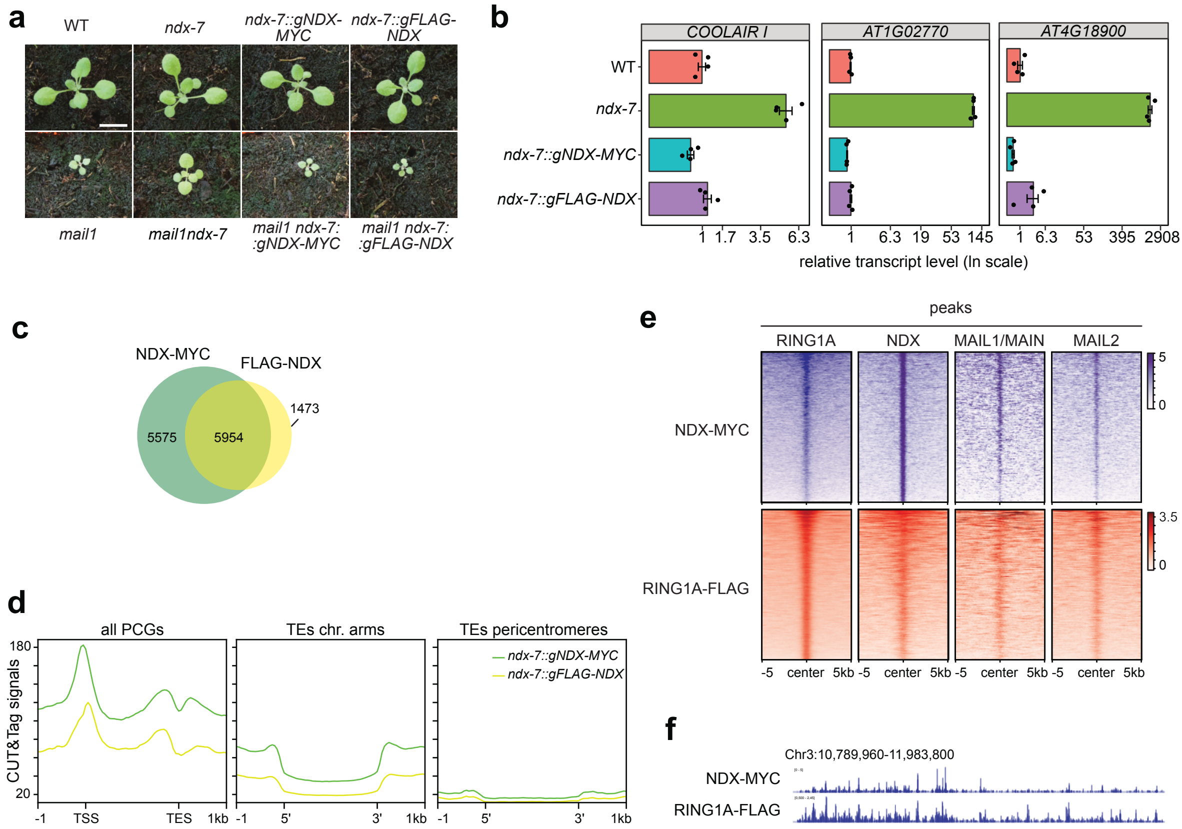


**Supplementary Fig. 8. NDX and RING1A are enriched at MAIL1/MAIN and MAIL2 binding sites** (**a**) Developmental phenotype of 2 week-old seedlings of indicated genotype. Scale bar is 1 cm. (**b**) Barplots illustrating molecular complementation of *ndx*-associated defects by tag-associated NDX transgene. Three loci up-regulated in the *ndx* mutant background are represented. Data are normalized to *ACT2* and further normalized to WT ; values represent means ± s.e.m computed from two technical RT-qPCR replicates on two biological replicates. (**c**) Venn diagrams illustrating the large overlap between NDX-MYC- and FLAG-NDX-associated peaks. (**d**) Metaplots showing NDX-MYC and FLAG-NDX signals in *ndx-7* complemented lines at protein-coding genes (left) and TEs located in chromosomal arms (middle) or pericentromeric regions (right). (**e**) Plot heatmaps showing enrichment of NDX and RING1A at NDX-myc/FLAG-NDX common peaks (n = 5954) and RING1A (n= 7706), MAIL1/MAIN (n = 626), MAIL2 (n = 1775) peaks. Peaks were aligned at their center position. (**f**) Genome browser view of a large chromosome 3 interval, illustrating the similarity between CUT&Tag NDX-MYC and ChIP-seq RING1A-FLAG genomic profiles.

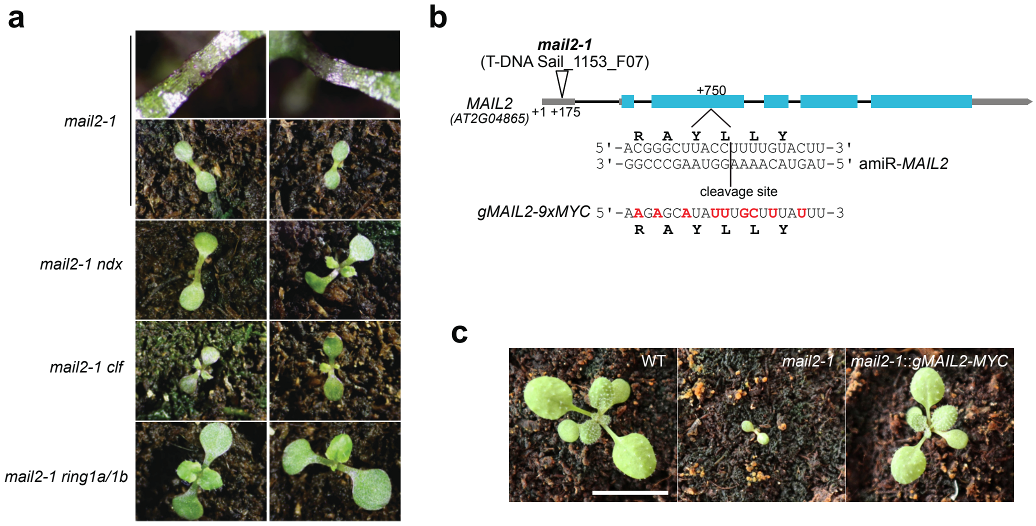


**Supplementary Fig. 9. *mail2-1* phenotypic defects are partially suppressed by *ring1a/1b* mutations.** (**a**) Developmental phenotypes associated to two-week-old seedlings of indicated genotype. (**b**) Schematic representation of *MAIL2* gene showing t-DNA *mail2-1* localization and amiR-*MAIL2* targeted sequence. Synonymous substitutions creating amiR-*MAIL2* insensitive *gMAIL2-9xMYC* transgene are indicated in red and corresponding amino acids are shown. Positions are given relative to the transcription start site (+1). (**c**) Two-week-old phenotype of WT, *mail2-1* and *gMAIL2-1-MYC* complemented *mail2-1* seedlings. Scale bar is 1 cm.

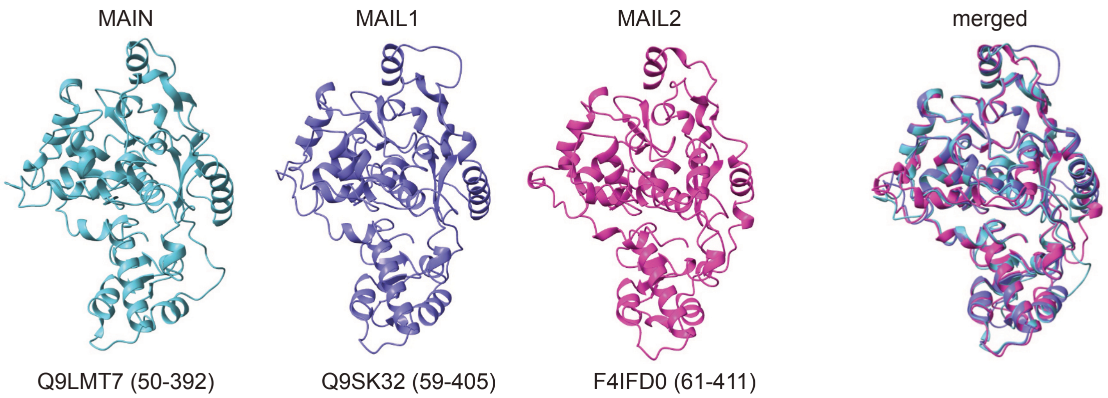


**Supplementary Fig. 10. MAIL1, MAIN and MAIL2 PMD-C domains displays highly similar 3D structures.** Structure predictions of MAIL1, MAIN and MAIL2 were recovered from the Alphafold Protein Structure Database and, individual and merged representation of PMD-specific parts are shown. UniProt IDs and position of PMD-C domains within the proteins are indicated below the structures.

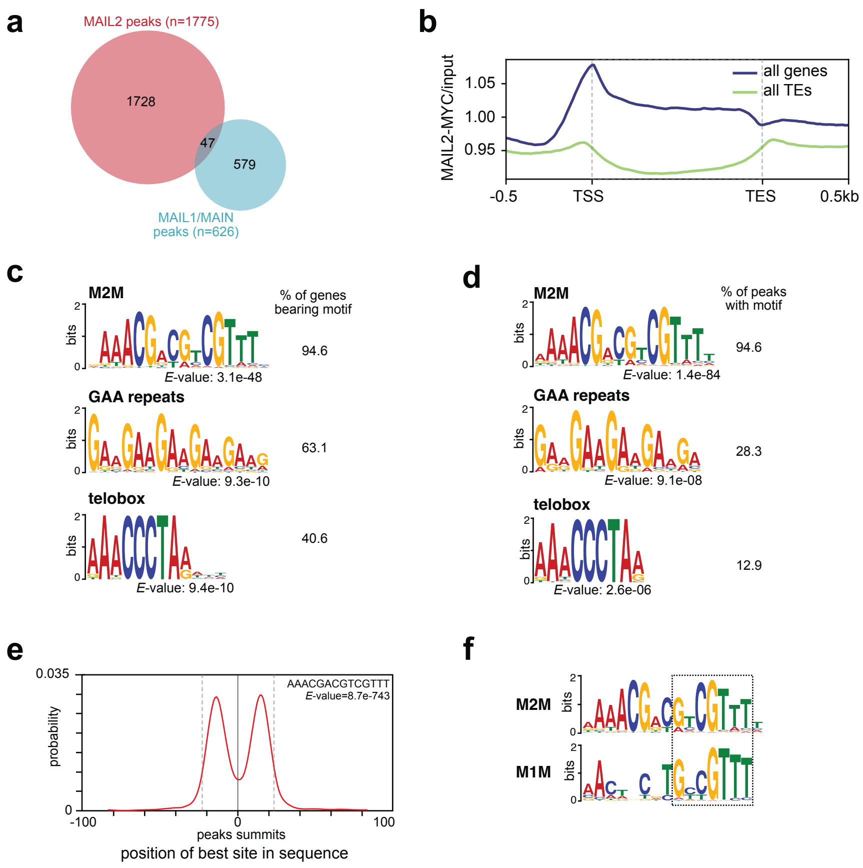


**Supplementary Fig. 11. MAIL2 targets a distinct set of genes through recognition of a specific DNA sequence motif.** (**a**) Overlap between MAIL1/MAIN and MAIL2-associated peaks. (**b**) Metagene plot showing mean MAIL2-MYC vs input signal at Arabidopsis gene and TE loci. (**c,d**) STREME prediction of conserved DNA motifs at MAIL2-associated gene promoters (n=1059) (**c**) or at MAIL2 peaks (n = 1775) (**d**). Relative representation of each motif is reported. (**e**) Centrimo plot showing strong enrichment of M2M over MAIL2 peaks, within a 45 bp central region delimited by the dashed lines (*E*-value = 8.7e-143). The 200bp region surrounding the peak summits is shown. (**f**) M1M homology to half of the palindromic M2M consensus sequence is highlighted by the dashed box.

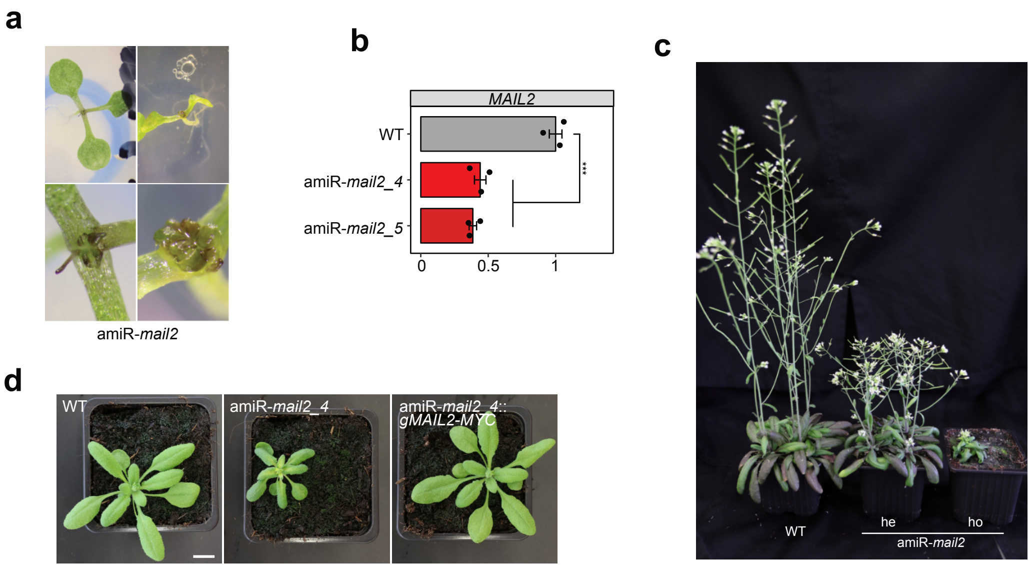


**Supplementary Fig. 12. Phenotype associated to amiR-*mail2* lines.** (**a**) Examples of developmental phenotype recovered from 10-day-old *in vitro* growing plants. (**b**) *MAIL2* transcript accumulation in WT and amiR-*mail2_4* and *_5* lines, detected by RT-qPCR using primers framing the amiRNA-targeted cutting site. Data are normalized to *ACT2* and further normalized to WT; values represent means from three biological replicates ± s.e.m and statistically significant differences are indicated (unpaired, two-sided Student's t-test, *** $P < 0.001$). (**c**) Photos of five-week-old WT plants compared to amiR-*mail2_4* mutant plants containing a single transgene locus at heterozygous (he) or homozygous (ho) state. (**d**) Two-week-old phenotypes of WT, amiR-*mail2* and *gMAIL2-1-9xMYC*-complemented amiR-*mail2* seedlings. Scale bar is 1 cm.

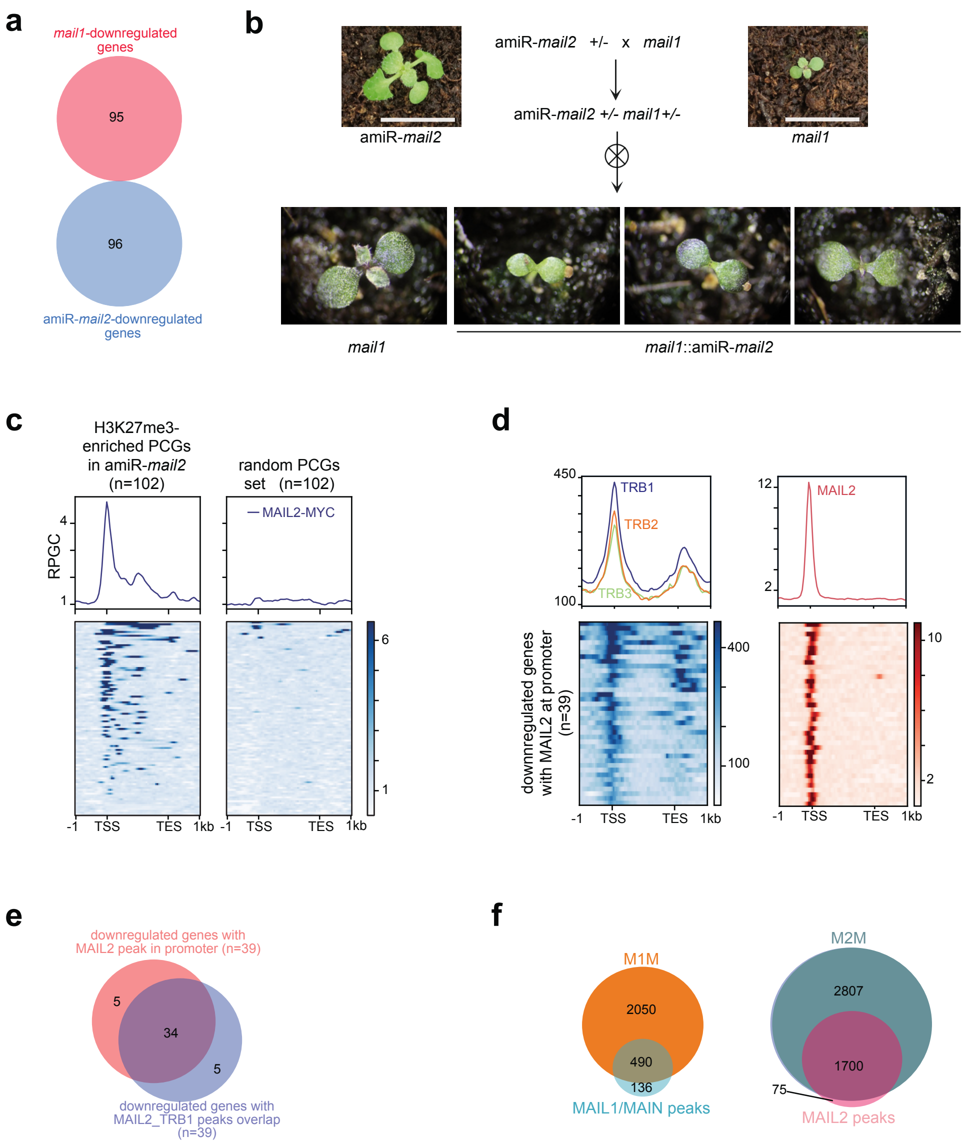


**Supplementary Fig. 13. MAIL1/MAIN and MAIL2 pathways target distinct sets of genes.** (**a**) Venn diagrams showing no overlap between *mail1*- and amiR-*mail2*- downregulated genes. (**b**) Developmental phenotypes observed upon MAIL2 depletion in *mail1* mutant background. Scheme of the cross is illustrated with photos of two-week-old plants. Scale bar is 1cm. Zoomed-in pictures of *mail1* F2 progeny expressing or not the amiR-*MAIL2* transgene are shown. (**c**) Metagene plots (top) and heatmaps (bottom) illustrating MAIL2 enrichment at PCGs that display enhanced H3K27me3 levels in amiR-*mail2* mutants (**d**) Metagene plots (top) and heatmaps (bottom) illustrating TRBs (data from[30]) and MAIL2 enrichment at MAIL2-associated promoter genes downregulated in amiRNA-*mail2* lines. For TRBs, only TRB1 data were represented on the heatmap. (**e**)Venn Diagram showing overlap between MAIL2 and TRB1 peaks at proximal promoter region of amiRNA-*mail2* downregulated genes. (**f**) Venn diagrams showing overlap of M1M and M2M loci with MAIL1 and MAIL2 peaks respectively.

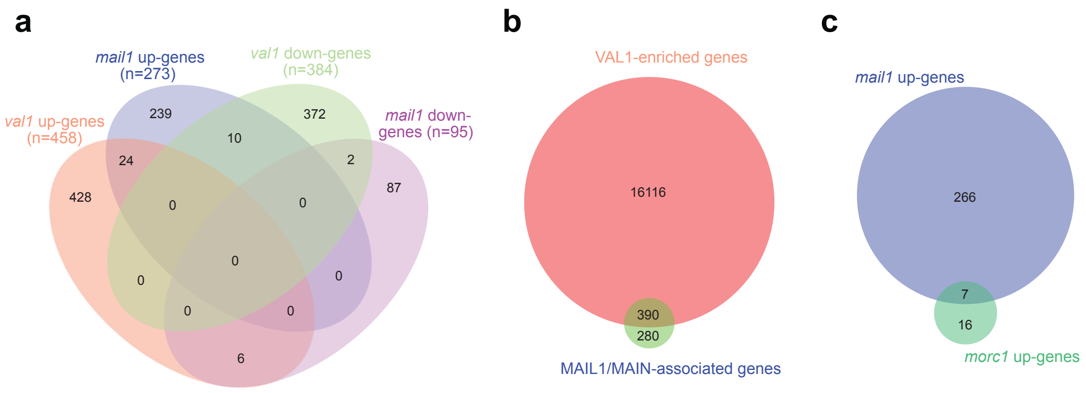


**Supplementary Fig. 14. VAL1 and MORC1 do not impact expression of *mail1* misregulated genes.** (**a,b**) Venn diagram illustration overlap (**a**) between *mail1* and *val1* misregulated genes or (**b**) between VAL1- and MAIL1/MAIN-associated genes. (**c**) Venn diagram showing overlap between *mail1*- and *morc1*-upregulated genes[23].

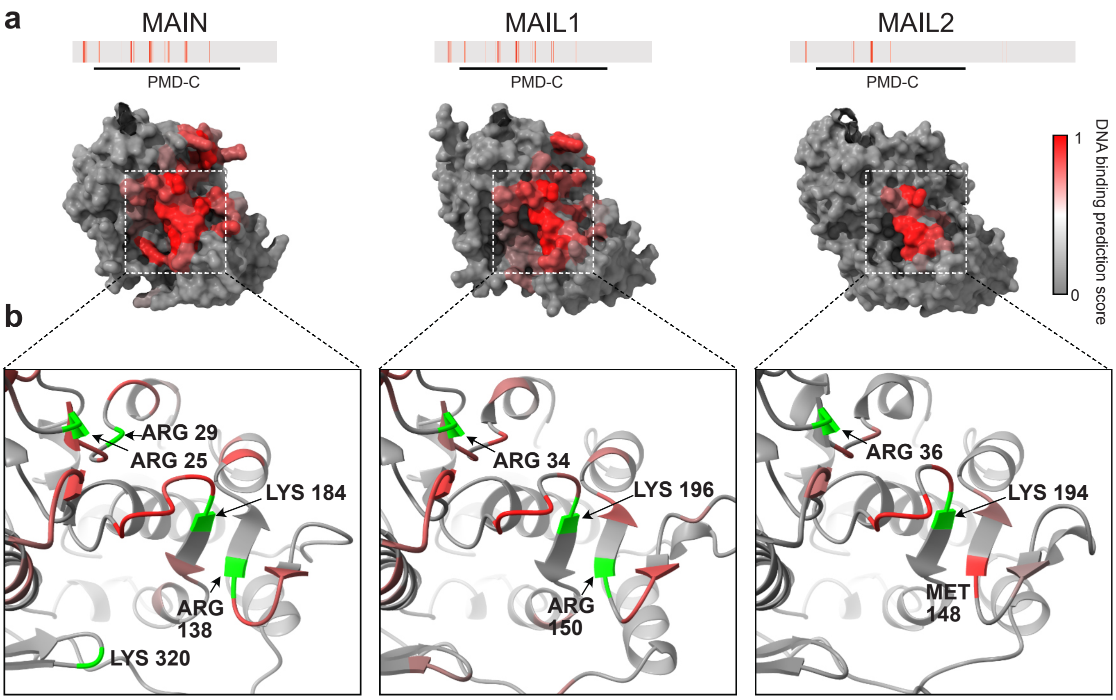


**Supplementary Fig. 15. Predicted DNA-binding potential of the PMD-C domains of MAIN, MAIL1 and MAIL2. (a)** Predicted structures of MAIN, MAIL1 and MAIL2 PMD-C domains, with DNA-binding residues mapped onto each structure and colored according to their prediction score. Full length proteins are shown above. **(b)** Close-up views of the indicated PMD-C regions. DNA-binding residues with a prediction confidence score ≥ 0.8 are colored as in (a). Positively charged residues with a confidence score ≥ 0.8 are highlighted in green. A complete list of DNA-binding prediction scores for amino acid residues of MAIN, MAIL1 and MAIL2 is reported in Supplementary Table 5.

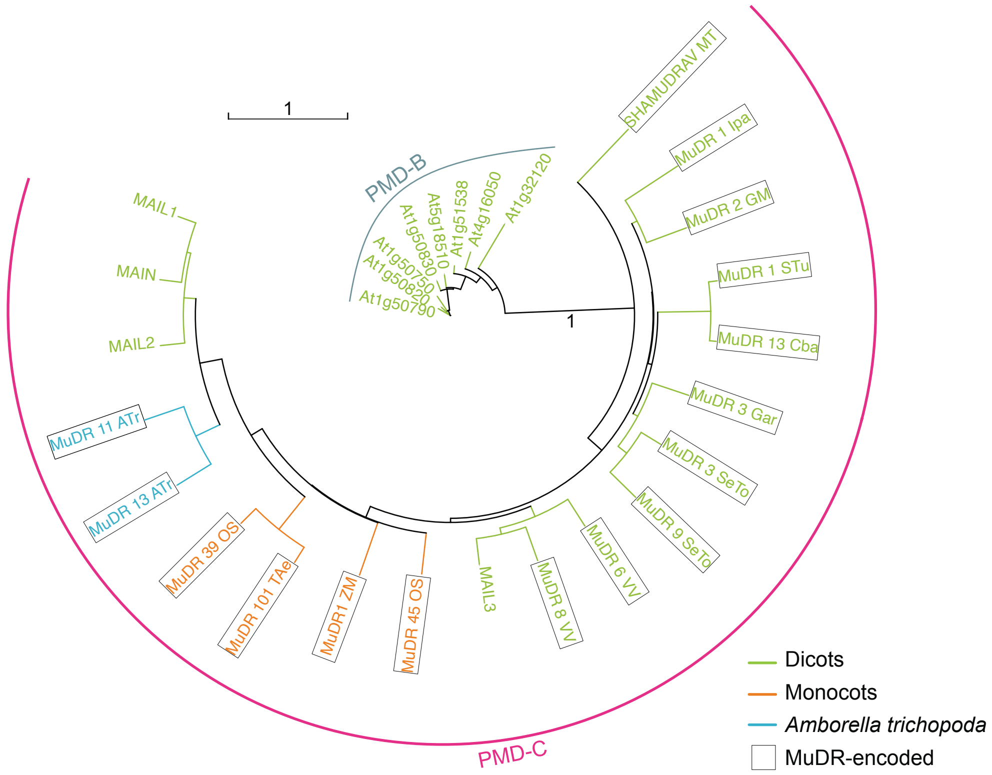


**Supplementary Fig. 16. PMD-C encoded by MuDR transposons are widely distributed in angiosperms**. A simplified phylogenetic tree was contructed using *Arabidopsis* gene-encoded PMD-B and PMD-C, and selected MuDR-encoded domains identified through tBlastN analysis of Repbase database[63] with MAIL2-specific PMD-C sequence as query (Supplementary doc. 1). Basal angiosperm *Amborella trichopoda*, monocot and dicot species are depicted in blue, orange and green, respectively. MuDR-encoded PMD-C were identified in *Amborella* and monocot species as previously reported[12], but were also found in dicotyledon species distributed across several Eudicots orders. Bootstrap value is indicated only for the node separating PMD-B and -C clades. Scale bar indicates one substitution/site. PMD sequences used to build the tree are reported in Supplementary Table 4. Abbreviations are: ATr *Amborella trichopoda*, Cba *Capsicum baccatum*, Gar *Gossypium arboretum*, GM *Glycine max*, Ipa *Ilex paraguariensis*, MT *Medicago truncatula*, OS *Oryza Sativa*, SeTo *Senna tora*, STu *Solanum tuberosum*, Tae *Triticum aestivum*, VV *Vitis Vinifera*, ZM *Zea mays*.
